# Robust reprogramming of glia into neurons by inhibition of Notch signaling and NFI factors in adult mammalian retina

**DOI:** 10.1101/2023.10.29.560483

**Authors:** Nguyet Le, Trieu-Duc Vu, Isabella Palazzo, Ritvik Pulya, Yehna Kim, Seth Blackshaw, Thanh Hoang

## Abstract

Generation of neurons through direct reprogramming has emerged as a promising therapeutic approach for neurodegenerative diseases. Despite successful applications *in vitro*, *in vivo* implementation has been hampered by low efficiency. In this study, we present a highly efficient strategy for reprogramming retinal glial cells into neurons by simultaneously inhibiting key negative regulators. By suppressing Notch signaling through the removal of its central mediator *Rbpj,* we induced mature Müller glial cells to reprogram into bipolar and amacrine neurons in uninjured adult mouse retinas, and observed that this effect was further enhanced by retinal injury. We found that specific loss of function of *Notch1* and *Notch2* receptors in Müller glia mimicked the effect of *Rbpj* deletion on Müller glia-derived neurogenesis. Integrated analysis of multiome (scRNA- and scATAC-seq) and CUT&Tag data revealed that Rbpj directly activates Notch effector genes and genes specific to mature Müller glia while also indirectly represses the expression of neurogenic bHLH factors. Furthermore, we found that combined loss of function of *Rbpj* and *Nfia/b/x* resulted in a robust conversion of nearly all Müller glia to neurons. Finally, we demonstrated that inducing Müller glial proliferation by AAV (adeno-associated virus)-mediated overexpression of dominant-active Yap supports efficient levels of Müller glia-derived neurogenesis in both *Rbpj*- and *Nfia/b/x/Rbpj*- deficient Müller glia. These findings demonstrate that, much like in zebrafish, Notch signaling actively represses neurogenic competence in mammalian Müller glia, and suggest that inhibition of Notch signaling and *Nfia/b/x* in combination with overexpression of activated Yap could serve as an effective component of regenerative therapies for degenerative retinal diseases.

## Introduction

Loss of neurons is the key pathological feature of many neurodegenerative diseases such as Parkinson’s disease, Alzheimer’s disease and retinal dystrophies. Photoreceptors and ganglion cells of the retina, like many other neuronal subtypes in the central nervous system, are susceptible to degeneration, with their loss often resulting in permanent blindness. While there is currently no effective regenerative therapy to replace neurons, one of the most potentially promising strategies is through direct reprogramming of endogenous Müller glia into retinal neurons. Müller glia are progenitor-like radial glial cells that serve as cellular sources for retinal regeneration in cold-blooded vertebrates. Zebrafish Müller glia respond to retinal injury by converting from a resting to an activated state, dividing asymmetrically to produce multipotent neurogenic retinal progenitor cells, and regenerating all types of retinal neurons ^1,2^. In mammals, however, Müller glia undergo reactive gliosis and do not spontaneously regenerate neurons following retinal injury.

Previous studies have shown that zebrafish Müller glia rapidly upregulate neurogenic bHLH factors such as *ascl1a* following injury ^3,4^. In mice, transgenic overexpression of *Ascl1* in combination with both histone deacetylase inhibition and other transcription factors (TFs) can induce Müller glia to reprogram into inner retinal-like neurons following injury ^5^. However, other factors act in parallel with these positive regulators of neurogenesis to repress neurogenic competence. Our previous work showed that the gene expression profile of resting mammalian Müller glia closely resembles retinal progenitor cells. These cells are robustly activated in response to injury, however, their conversion to a neurogenic state is actively inhibited by a dedicated transcriptional regulatory network that restores Müller glia to a resting state ^6^.

Within this network, we identified the Nuclear Factor I (NFI) family of TFs as negative regulators of injury-induced reprogramming. Simultaneous loss of function of *Nfia, Nfib,* and *Nfix (Nfia/b/x)* in mature mouse Müller glia lead a subset of cells to convert to bipolar or amacrine cells primarily through a process of direct transdifferentiation ^6^, much like what has been observed following *Ascl1* overexpression ^5^. Therefore, combinatorial approaches that target both positive and negative regulators of neurogenic competence in Müller glia are likely needed for the development of effective reprogramming therapies.

However, major gaps in our understanding of the regulation of neurogenic competence in Müller glia must be bridged before therapeutic applications can be further developed. Current methods for inducing neurogenic competence, whether involving bHLH overexpression or NFI factors loss of function often require retinal injury to induce Müller glia reprogramming ^5,6^. Furthermore, the levels of induced neurogenesis are relatively low, and the modest levels of Müller glia proliferation raise the concerns about potential depletion of Müller glia if the efficiency of neurogenesis is enhanced. Moreover, the range of cell types generated from Müller glia is typically limited to bipolar and amacrine cells, although recent studies have also successfully generated retinal ganglion-like cells ^7,8^. Therefore, it is imperative to conduct further research aiming at identifying additional candidate regulators, both positive and negative, of neurogenic competence to advance our understanding and develop more effective strategies for efficient glia-to-neuron reprogramming in the retina.

Our previous work has identified transcriptional effectors of the Notch pathway as key candidate regulators of Müller glial quiescence in both zebrafish and mice ^6^. Notch signaling plays a central role in controlling vertebrate retinal development, where it maintains progenitor status, promotes gliogenesis and inhibits neurogenesis, particularly photoreceptor specification ^9–13^. In mature zebrafish Müller glia, it actively represses injury-induced reprogramming ^14,15^. Suppression of Notch signaling via gamma-secretase inhibitors efficiently induces injury-independent reprogramming and enhances injury-induced reprogramming ^16–20^. In mammals, low levels of Notch signaling are detected in mature Müller glia ^6,16^, but it remains unclear whether active Notch signaling acts to maintain Müller glia quiescence and represses neurogenic competence in these cells.

In this study, we assessed whether loss of Notch signaling could induce reprogramming of mammalian Müller glial cells. We found that selective deletion of the common Notch transcriptional mediator *Rbpj* in adult mouse Müller glia induced direct transdifferentiation into bipolar and amacrine cells without injury, and that this process was significantly enhanced by NMDA-dependent excitotoxic injury. Furthermore, Müller glia-specific loss of function of *Notch1* and *Notch2* receptors phenocopied the effect of *Rbpj* deletion. We also showed that simultaneous loss of function of *Rbpj* and *Nfia/b/x* led to a conversion of nearly all Müller glia to neurons and that AAV-mediated overexpression of dominant-active Yap induced these cells to proliferate. These findings demonstrate that, much like in zebrafish, Notch signaling actively represses neurogenic competence in mammalian Müller glia, and imply that efficient endogenous cell reprogramming can be achieved by targeting both negative and positive regulators of neurogenesis.

## Results

### Suppression of RBPJ-mediated Notch signaling induces MG-derived neurogenesis in the adult mouse retina with and without retinal injury

To comprehensively examine the cellular expression patterns of Notch pathway components in adult retina, we analyzed scRNA-Seq datasets obtained from mice and humans ^6,21,22^. Our scRNA-Seq data analysis showed an evolutionary conserved expression of Notch signaling components in both mouse and human Müller glia. In both species, Müller glia expressed *Notch1* and *Notch2*, and substantially lower levels of *Notch3* and *Notch4* (Fig. S1A-B). Expression of all key Notch ligands, mediators and target genes, including *Jag1*, *Rbpj*, *Numb*, *Hes1*, *Hes5*, *Hey1*, *Hey2* and *Id1/2/3/4*, were found in both mouse and human Müller glia. No other retinal cell type, except vascular/endothelial cells, showed comparable expression of Notch signaling components.

Previous studies showed that loss of function of the common Notch transcriptional mediator *Rbpj* resulted in global inhibition of Notch signaling ^23–25^ and induced a limited level of astrocyte-to-neuron in brain conversion following injury ^26–28^. To assess whether loss of Notch signaling could induce reprogramming of mature mouse Müller glia, we selectively deleted *Rbpj* in Müller glia using *GlastCreER*^T2^;*Rbpj^lox/lox^;Sun1-GFP* mice (Fig. 1A). These transgenic mice express a tamoxifen-inducible Cre recombinase under the control of regulatory elements of the Müller glia-specific Glast (*Slc1a3*) gene. We induced Cre recombination in *GlastCreER*^T2^;*Sun1-GFP* control and *GlastCreER*^T2^;*Rbpj^lox/lox^;Sun1-GFP* mice with five daily intraperitoneal (i.p.) injections of tamoxifen. Tamoxifen administration leads to both Cre-dependent removal of exon 4 of *Rbpj* that results in a functional null mutation ^29^ and expression of *Sun1- GFP* reporter, thereby allowing lineage tracing of all Müller glia and Müller glia-derived cells ^6,30^. Uninjured retinas were collected for immunohistochemistry (IHC) analysis at multiple timepoints ranging from 3 weeks to 4 months post tamoxifen injection (Fig. 1B). In previous studies, retinal injury and associated glial activation were required for inducing Müller glia-derived neurogenesis following either *Ascl1* overexpression or *Nfia/b/x* loss of function ^5,6^. We also tested whether N-methyl-d-aspartate (NMDA)- mediated excitotoxic injury, which primarily causes cell death of amacrine and retinal ganglion cells, could enhance neurogenesis in *Rbp*j-deficient Müller glia. Intravitreal NMDA injection was performed at 7 days after last dose of tamoxifen and retinas were analyzed at 3 weeks, 5 weeks and 4 months following NMDA injury (Fig. 1C).

**Figure 1:**
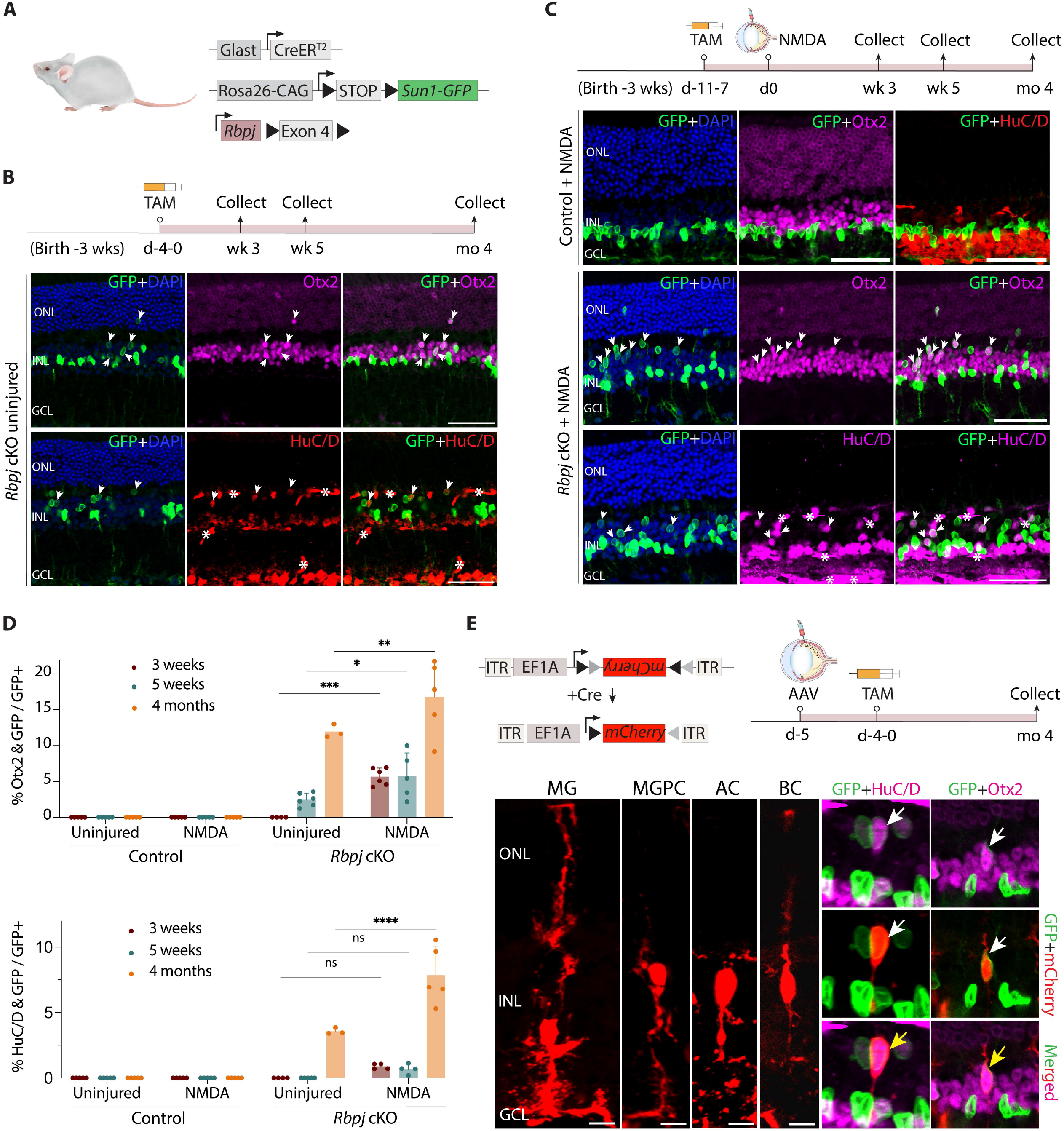
Selective loss of function of *Rbpj* induces neurogenesis in adult Müller glia with and without retinal injury. **(A)** Schematic of the transgenic constructs used to induce deletion of *Rbpj* specifically in MG. Cre-mediated removal of Exon 4 of *Rbpj* leads to premature stop codon formation and disrupts the expression of the RBPJ protein. **(B,C)** Schematic of the experimental workflow and representative immunostaining for GFP, Otx2, and HuC/D in control and *Rbpj*-deleted retinas without (**B**) and with (**C**) NMDA-induced injury. White arrowheads indicate co-labeled GFP-positive & marker-positive cells. *Asterisks indicate mouse-on-mouse vascular staining. Scale bar = 50μm. **(D)** Quantification of mean percentage ± SD of GFP-positive MG-derived neurons expressing either OTX2 or HuC/D. **(E)** Morphological characterization of MG-derived neurons in *Rbpj*-deficient retinas using AAV2.7m8-Ef1a- Flex-mCherry 4 months post NMDA treatment. Yellow arrowheads indicate co-labeled GFP/mCherry/marker-positive cells. Scale bar = 10μm. Significance was determined via two-way ANOVA with Tukey’s multiple comparison test: ****p < 0.0001. Each data point was calculated from an individual retina. TAM, tamoxifen; ONL, outer nuclear layer; INL, inner nuclear layer; GCL, ganglion cell layer, MG, Müller glia; MGPC, Müller glia-derived progenitor cell; AC, amacrine cell; BC, bipolar cell.

We detected no Müller glia-derived neurogenesis in the uninjured and NMDA-treated control retinas and all GFP-positive cells expressed the glial marker Sox9 (Fig. 1C, S2A). However, in *Rbpj* conditional mutants, we observed that 2.5% of GFP-positive Müller glia-derived cells expressed the pan-bipolar/photoreceptor cell marker Otx2 at 5 weeks, with a progressive increase to 12% at 4 months post-induction (Fig. 1B, D). At this later timepoint, we also observed 3.6% of GFP-positive cells expressing the pan-amacrine/retinal ganglion cell marker HuC/D (Fig. 1B, D). NMDA treatment substantially enhanced levels of Müller glia-derived neurogenesis, with 5.8% and 15% of GFP-positive cells expressing Otx2 at 5 weeks and 4 months respectively (Fig. 1C, D). The levels of Müller glia-derived amacrine cells did not change significantly at 3 and 5 weeks in NMDA-treated retinas compared to uninjured retinas. However, they were enhanced at 4 months, with 7.8% of GFP-positive cells in showing HuC/D expression, in contrast to 3.6% of GFP-positive cells in uninjured retina (Fig. 1C, D).

IHC analysis of additional cell markers revealed a subset of *Rbpj*-deficient Müller glia-derived neurons expressed the glycinergic amacrine-specific marker Glycine (Fig. S2C) and the cone bipolar-specific marker Scgn (Fig. S2D), but did not express rod bipolar markers such as PKCa (Fig. S2E). No GFP-positive cells expressing rod photoreceptor-specific markers such as Nrl or retinal ganglion cells such as Rbpms were observed and most GFP-positive cells were confined to the inner nuclear layer (INL). In addition to neurogenesis, we also examined whether loss of *Rbpj* induces Müller glia glia-derived proliferation by injecting 7 daily doses of 5-ethynyl-2’-deoxyuridine (EdU) at 1 week post tamoxifen. EdU labeling revealed no GFP-positive cells incorporating EdU, nor was Ki67 staining observed, indicating that *Rbpj*-deficient Müller glia did not undergo proliferation (Fig. S2F). Expression of neurogenic bHLH factors such as *Ascl1* was activated in *Rbpj*-deficient Müller glia (Fig. S2G). Sox9 expression was detected in the remaining GFP-positive cells that did not undergo neurogenesis, indicating that they retained glial identity (Fig. S2H).

The use of nuclear membrane-bound Sun1-GFP reporter is advantageous for unambiguous identification of Müller glia-derived neurons, however the overall cell morphology cannot be observed. To better characterize the neuronal morphology of newborn neurons, we performed intravitreal injection of Cre-dependent FLEX AAV carrying mCherry reporter into adult *Rbpj* conditional knockout mice. We then induce Cre recombination with 5 daily i.p. injections of tamoxifen and harvested the retinas 4 months later. IHC analysis revealed that the newborn neurons retracted their apical glial processes and acquired bipolar or amacrine-like morphologies, indicated by coexpression of mCherry/GFP with either Otx2 or HuC/D (Fig. 1E). Together, these findings indicate that Notch signaling negatively regulates Müller glia neurogenic competence in mammals and suppressing this pathway induces Müller glia to transdifferentiate into bipolar and amacrine cells without injury and this process is significantly enhanced with injury.

### Loss of function of *Notch1/2* receptors phenocopies neurogenesis observed in *Rbpj*- deficient Müller glia

While *Rbpj* is a well-established transcriptional mediator of Notch signaling, previous studies suggest that it may have Notch-independent functions ^31–33^, and conversely that some aspects of Notch signaling may be *Rbpj*-independent ^34,35^. There are 4 Notch receptors, Notch1-4, with Notch1 and Notch2 being the two most abundantly expressed in mature Müller glia (Fig. S1). To assess whether repression of neurogenic competence in Müller glia was indeed dependent on active Notch signaling, we generated *GlastCreER*^T2^;*Notch1^lox/lox^;Sun1-GFP*, *GlastCreER*^T2^;*Notch2^lox/lox^;Sun1-GFP* single and *GlastCreER*^T2^;*Notch1^lox/lox^;Notch2^lox/lox^;Sun1-GFP* double loss of function mutants of *Notch1* and *Notch2* ^36,37^ (Fig. 2A). As previously described, we performed 5 daily i.p. injections of tamoxifen to delete *Notch1* and/or *Notch2* specifically in Müller glia while simultaneously labeling these cells with Sun1-GFP for lineage tracing (Fig. 2B). We then induced excitotoxic retinal injury with NMDA and analyzed the retina 5 weeks later. We did not observe any induction of neurogenesis in *Notch1*-deficient Müller glia, and observed that only 0.7% of GFP-positive cells expressed Otx2 in *Notch2*-deficient mice. However, *Notch1/2* double mutant retinas showed similar levels of Müller glia-derived bipolar cells seen in age-matched *Rbpj*-deficient retinas in the absence of retinal injury (Fig. 2C,D). The relative numbers of Müller glia-derived HuC/D-positive amacrine cells, however, were significantly lower in *Notch1/2* double mutants relative to *Rbpj* mutants in the absence of injury (Fig. 2D). Likewise, the level of Müller glia-derived neurogenesis in *Notch1/2* double mutants is also comparable to those seen with *Rbpj* mutants following NMDA injury (Fig. 2E-G). As in *Rpbj* conditional knockout mice, a subset of *Notch1/2*-deficient Müller glia- derived neurons also expressed the cone bipolar-specific marker Scgn (Fig. S2I), but did not express rod bipolar markers such as PKCa (Fig. S2J). Expression of neurogenic bHLH factors such as *Ascl1* was activated in *Notch1/2*-deficient Müller glia (Fig. S2K) and Sox9 expression was detected in the remaining GFP-positive cells, indicating that they retained glial identity (Fig. S2L). These results demonstrate that *Notch1/2* loss of function effectively phenocopies *Rbpj* loss of function.

**Figure 2:**
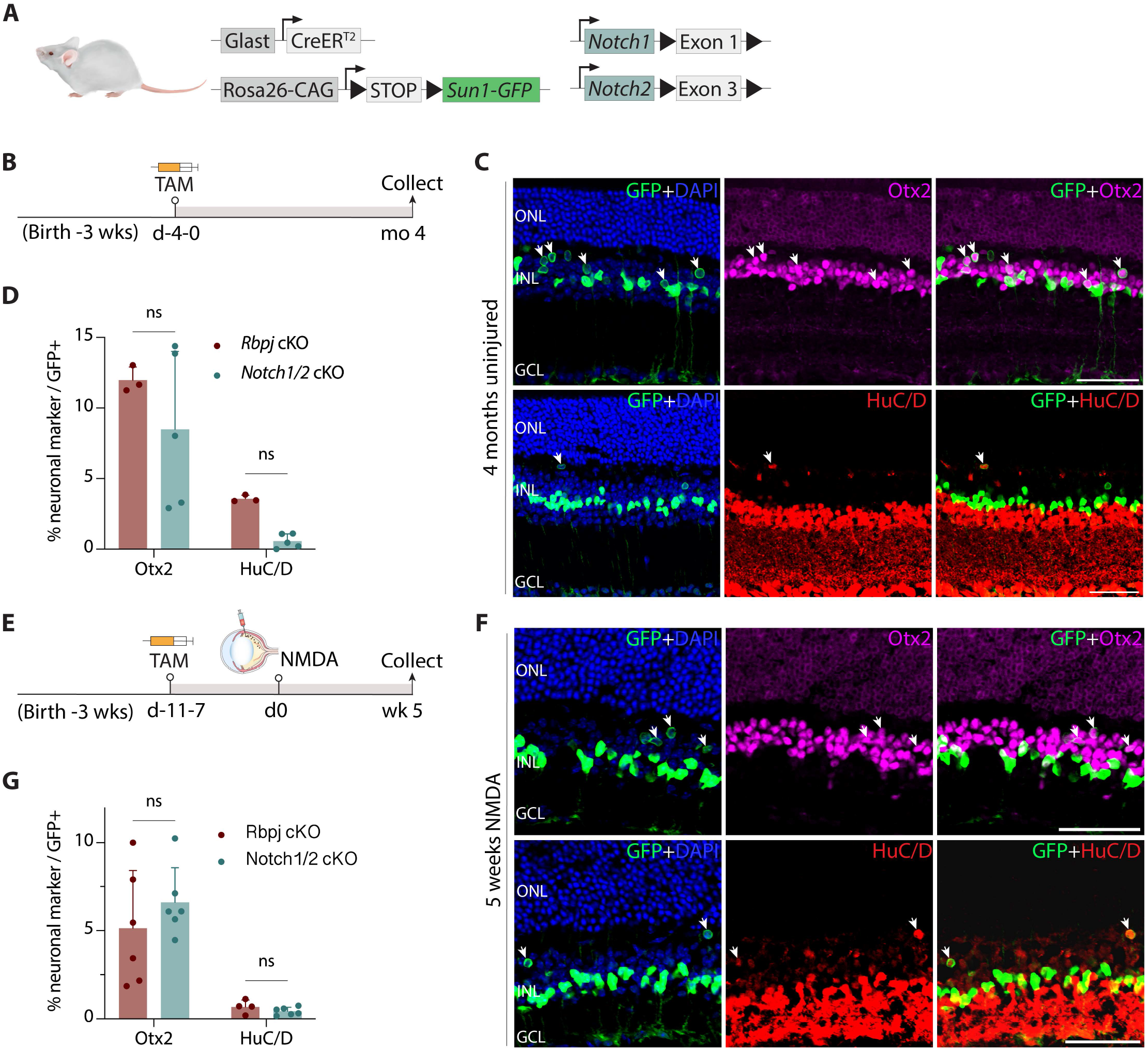
*Notch1/2* deletion phenocopies effects of *Rbpj* deletion on Müller glia-derived neurogenesis. **(A)** Schematic of the transgenic constructs used to induce deletion of *Notch1 and Notch2* specifically in MG. Cre-mediated removal of Exon 1 of *Notch1* and Exon 3 of *Notch2* leads to premature stop codon formation and disrupts the expression of the NOTCH1/2 proteins. **(B)** Schematic of the experimental workflow for uninjured retinas. **(C,D)** Representative images immunolabeled for GFP, OTX2, and HuC/D of Notch1/Notch2-deleted retinas after 4 months post tamoxifen injection, and quantification of mean percentage ± SD of GFP-positive MG-derived neurons expressing either Otx2 or HuC/D. White arrowheads indicate co-labeled GFP-positive & marker-positive cells. **(E-G)** Schematic of the workflow, representative immunostaining, and quantification for NMDA-injured retinas. Significance was determined via Mann-Whitney U test: ****p < 0.0001. Each data point was calculated from an individual retina. TAM, tamoxifen; ONL, outer nuclear layer; INL, inner nuclear layer; GCL, ganglion cell layer. Scale bar = 50μm.

### Rbpj directly promotes expression of Notch pathway genes and genes specific to resting Müller glia, while indirectly repressing neurogenic bHLH factors

To further investigate the effects of *Rbpj* loss of function, we conducted scRNA-Seq on uninjured whole retinas of *GlastCreER*^T2^;*Sun1-GFP* control and *GlastCreER*^T2^;*Rbpj^lox/lox^;Sun1-GFP* mice at 7 days following tamoxifen injection (Fig. 3A, Fig. S3A). We then subsetted the Müller glia population for further downstream analysis. At 7 days post tamoxifen injection, before any neurogenesis is detected, while Müller glia from both samples show broadly similar transcriptional profiles (Fig. 3B), a subset of genes show strongly enriched expression in *Rbpj*-deficient Müller glia. Genes specific to mature Müller glia (*Glul*, *Aqp4*, *Apoe*, *Kcnj10*, *Mlc1*, *Sox9*) and Notch pathway genes (*Hes1*, *Hes5*, *Id1/2/3*) are significantly downregulated in *Rbpj*-deficient Müller glia relative to controls. In contrast, neurogenic bHLH factors (*Ascl1*, *Neurog2*, *Hes6*) are strongly upregulated (Fig. 3C,D, Fig. S3B,C, Table ST1). Interestingly, expression of cell cycle inhibitors such as *Cdkn1b, Cdkn1c* and *Btg2*, is upregulated, consistent with the observation that *Rbpj*- deficient Müller glia undergo direct conversion into retinal neurons without proliferation.

**Figure 3:**
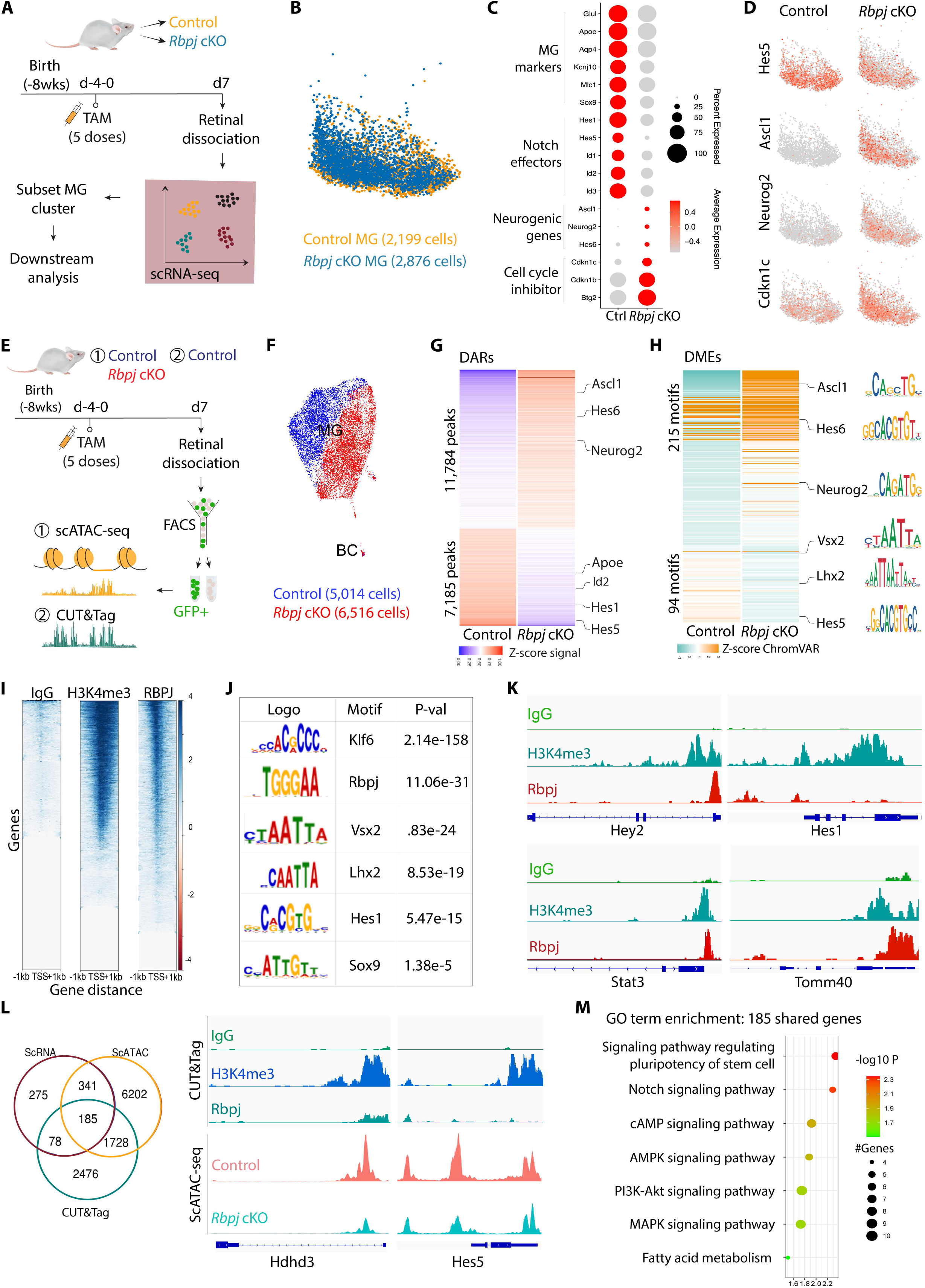
*Rbpj* deletion in Müller glia downregulates Notch target genes and activates expression of neurogenic genes. **(A)** Schematic of the experimental workflow used to generate scRNA-Seq data from whole retinas of control and *Rbpj* conditional knockout mice. **(B)** UMAP of scRNA-Seq data showing the clustering of control and *Rbpj*-deficient MG subsetted from whole retina cell populations. **(C)** Dot plot showing downregulation of genes specific to resting MG and Notch-regulated genes, and upregulation of neurogenic bHLH factors as well as cell cycle inhibitors in *Rbpj*-deficient MG relative to controls. **(D)** Feature plots highlighting differential expression of *Hes5*, *Ascl1*, *Neurog2*, and *Cdkn1c* in control and *Rbpj*- deficient MG. **(E)** Schematic of the experimental workflow used to generate scATAC-seq data from control and *Rbpj* cKO MG and CUT&Tag data from control MG. **(F)** UMAP plot showing global differences in chromatin accessibility observed by scATAC-Seq in control and *Rbpj*-deficient MG. **(G)** Differential accessibility regions (DARs) and nearby genes between *Rbpj*-deficient and control Müller glia. **(H)** Differential ranscription factor motif enrichments (DMEs) in the DARs between *Rbpj*-deficient and control Müller glia. **(I)** CUT&Tag analysis of IgG, H3K4me3, and RBPJ from FACS-isolated GFP-positive Müller glia from *GlastCreER^T^*^2^*;Sun1GFP* retinas. **(J)** Transcription factor motifs enriched in RBPJ CUT&Tag samples relative to control. **(K)** Representative genome tracks for CUT&Tag analysis. **(L)** Integrative analysis of scRNA-Seq, scATAC-Seq, and CUT&Tag to identify genes regulated by RBPJ. **(M)** Gene Ontology enrichment analysis for genes shared among the three datasets.

To examine the changes in chromatin accessibility caused by *Rbpj* deletion, we performed scATAC-seq on FACS-isolated GFP-positive cells from uninjured retinas of *GlastCreER*^T2^;*Sun1-GFP* control and *GlastCreER*^T2^;*Rbpj^lox/lox^;Sun1-GFP* mice at 7 days following tamoxifen injection (Fig. 3E). Clustering analysis showed a much clearer separation of control and *Rbpj*-deficient Müller glia cells than was seen using scRNA-Seq, indicating that loss of function of *Rbpj* leads to global changes in chromatin accessibility (Fig. 3F, Fig. S3E). We then compared control and *Rbpj*-deleted Müller glia, and identified 18,969 differential accessible chromatin regions (DARs) between the two samples. Furthermore, we also observed that changes in chromatin accessibility broadly mirror changes in transcription, with decreased accessibility at Notch pathway genes such as *Hes1*, *Hes5, Hey2* and *Id1/2/3/4*, and increased accessibility at neurogenic bHLH genes such as *Ascl1, Neurog2* and *Hes6* (Fig. 3G, Table ST1). We next examined TF motif enrichment in the DARs regions. As expected, we observed significant motif enrichments for neurogenic TFs, including *Ascl1, Neurog2* and *Hes6,* and reduction in accessibility associated with TF motifs for Müller glia-enriched genes, such as *Rbpj*, *Lhx2*, *Hes5* and *Vsx2,* in the *Rbpj*-deficient Müller glia (Fig. 3H, Fig. S3D).

To identify genomic binding targets of Rbpj, we next performed CUT&Tag on FACS-isolated GFP- positive adult Müller glia from *GlastCreER*^T2^;*Sun1-GFP* mice at 7 days following tamoxifen injection (Fig. 3E,I). Rbpj-binding genomic regions were significantly enriched for consensus motifs for Rbpj and Müller glia-enriched TFs such as Klf6, Lhx2, Hes1, Hes5 and Sox8/9 (Fig. 3J). As expected, Rbpj binds directly to cis-regulatory sites associated with Notch pathway genes, including *Hes1*, *Hes5*, *Hey2* and *Id1/2/3/4* as well as other TFs such as *Tcf7l2* that are known to promote gliogenesis^38^ (Fig. 3K, Fig. S3F, Table ST2).

We found that Rbpj also directly binds to regulatory sites associated with genes that promote cell cycle inhibition, including *Cdkn1c* and *Btg2* (Table ST2). We next sought to comprehensively identify genes directly regulated by Rbpj by integrating differential gene expression, DARs and Rbpj-binding regions from scRNA-Seq, scATAC-Seq and CUT&Tag data, respectively. We found that 185 genes were shared among the three datasets, indicating the expression of these genes was directly controlled by Rbpj (Fig. 3L). Gene Ontology (GO) enrichment analysis showed that these genes were enriched for Notch, Akt, and MAPK signaling, as well as fatty acid metabolism (Fig. 3M). Together, our findings revealed that Rbpj directly activates Notch effector genes and genes specific to mature Müller glia while also indirectly represses the expression of neurogenic bHLH factors.

### Combined loss of function of *Nfia/b/x* and *Rbpj* leads to conversion of the majority of Müller glia to neurons

We previously showed that the NFI factors inhibit neurogenic competence in mature Müller glia ^6^. Additional analysis of *Nfia/b/x*-deficient Müller glia demonstrates that Notch pathway genes such as *Hes1*, *Hes5*, and *Hey2* remained expressed in *Nfia/b/x*-deficient Müller glia (Fig. S4A-C). Furthermore, the expression and motif activity of *Nfia/b/x* were largely unaffected in *Rbpj*-deficient Müller glia (Fig. S3C-D, Table ST1). This raises the question whether Notch signaling and NFI factors act in parallel to inhibit neurogenic competence in adult Müller glia. To address this, we used the gamma-secretase inhibitor DAPT to inhibit Notch signaling and our previously described Müller glia-specific tamoxifen inducible mouse model of *Nfia/b/x* loss of function (*GlastCreER*^T2^*;Nfia/b/x^lox/lox^; Sun1-GFP*)^6^. Briefly, adult mice were fed tamoxifen diet for 3 weeks, followed by intravitreal injection of DAPT, then NMDA, and a second dose of DAPT (Fig. S4D). Retinas were then collected 3 weeks later for immunostaining analysis. Quantification of HuC/D/GFP-positive cells revealed over 2-fold increase in the number of Müller glia-derived neurons in DAPT-treated retinas compared to vehicle control (Fig. S4D). This finding strongly supports our hypothesis that *Nfia/b/x* and *Rbpj* act in parallel to repress Müller glia neurogenic competence.

To better assess potential combinatorial role for *Nfia/b/x* and *Rbpj* in repressing neurogenic competence, we generated *GlastCreER*^T2^;*Nfia/b/x^lox/lox^;Rbpj^lox/lox^ ;Sun1-GFP* mice (Fig. 4A). This transgenic mouse model allows a more specific inhibition of the Notch signaling pathway in Müller glia than global inhibition of gamma-secretase, and also serves as a cross-validation of the DAPT treatment. After 5 daily i.p. injections of tamoxifen to delete *Nfia/b/x* and *Rbpj* in Müller glia, we induced retinal injury with injection of NMDA in the right eye while leaving the left eye uninjured. We then harvested the retinas for immunostaining analysis 3 weeks later to examine the level of Müller glia-derived neurogenesis (Fig. 4B). We observed that combined loss of function of *Nfia/b/x* and *Rbpj* leads to a robust and synergistic increase in neurogenesis relative to both *Nfia/b/x*- ^6^ and *Rbpj-*deficient Müller glia (Fig. 1). In uninjured *Nfia/b/x;Rbpj*- deficient retinas, we found that 45.3% of GFP-positive cells expressed either Otx2 or HuC/D at 4 weeks following tamoxifen induction, with many of these cells undergoing radial migration into the outer nuclear layer (Fig. 4C,D).

**Figure 4:**
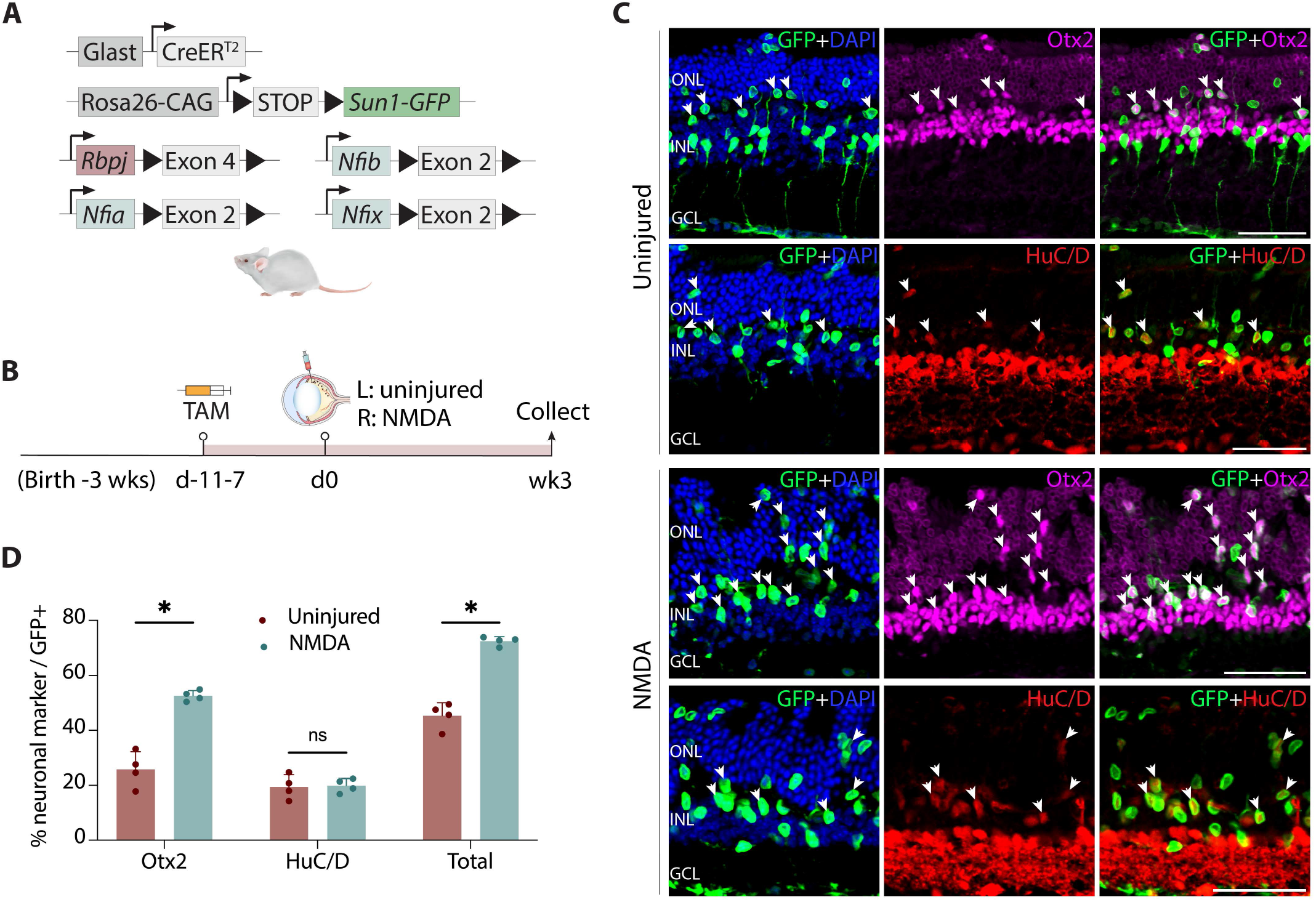
Simultaneous disruption of *Nfia/b/x* and *Rbpj* convert nearly all adult Müller glia into retinal neurons. **(A)** Schematic of the transgenic constructs used to induce loss of function of *Rbpj* and *Nfi/a/b/x* specifically in Müller glia. **(B,C)** Schematic of experimental pipeline and representative images of uninjured and NMDA-damaged retinas immunolabeled for GFP, Otx2 and HuC/D. White arrowheads indicate GFP-positive MG-derived neurons expressing neuronal markers Otx2 or HuC/D. **(D)** Quantification of mean percentage ± SD of GFP-positive Müller glia-derived neurons expressing either Otx2 or HuC/D in uninjured and NMDA-treated retina. Significance was determined via unpaired t test: ****p < 0.0001. TAM, tamoxifen; ONL, outer nuclear layer; INL, inner nuclear layer; GCL, ganglion cell layer. Scale bar = 50μm.

NMDA-mediated excitotoxic injury induces even higher levels of glia-to-neuron conversion in *Nfia/b/x;Rbpj*-deficient Müller glia. At 3 weeks post-injury, we observe that the vast majority of Sun1-GFP- positive cell nuclei are radially displaced away from their typical location in the INL, with many now present in the outer nuclear layer (ONL) (Fig. 4C). We observe that 52.6% of all GFP-positive Müller glia-derived cells are strongly immunopositive for Otx2, indicating that they are likely bipolar cells, while 19.9% are HuC/D-positive amacrine cells (Fig. 4C,D). Similar to what is seen in *Rpbj*-deficient retinas, a subset of *Nfia/b/x;Rbpj*-deficient Müller glia-derived neurons also expressed the cone bipolar-specific marker Scgn (Fig. S5A), but did not express the rod bipolar marker PKCa (Fig. S5B). No EdU incorporation was detected (Fig. S5C). We also observe a significantly reduced number of GFP-positive cells expressing Sox9 in both uninjured and NMDA-treated retina relative to wildtype controls (Fig. S5D). In line with this massive reduction in the number of Müller glia due to direct transdifferentiation, we also observe widespread disruptions in retinal lamination, with whorls and rosettes formed throughout the ONL (Fig. 4C). This disorganization of the outer limiting membrane mimics the effects observed by selective loss of a large fraction of Müller glia ^39–42^. In addition to these Müller glia-derived bipolar and amacrine, we also observed small numbers of GFP-positive cells expressing Nrl or Recoverin in the ONL, indicating that we also are generating some rod-like photoreceptors (Fig. S5E,F).

To better understand the molecular mechanism underlying Müller glia reprogramming in the context of *Nfia/b/x* and *Rbpj* deficiency, we conducted single nuclei multi-omic (combined RNA- and ATAC-seq) analysis of FACS-isolated GFP-positive cells from control *GlastCreER*^T2^*;Sun1-GFP* and *GlastCreER*^T2^*;Nfia/b/x^lox/lox^; Sun1-GFP* mice. Both uninjured and NMDA-treated retinas were profiled 4 weeks following tamoxifen induction (Fig. 5A, Table ST3). The combined UMAP plot of the scRNA-seq and scATAC-seq data show a clear separation of control and *Nfia/b/x;Rbpj*-deficient Müller glia clusters, suggesting that deleting *Nfia/b/x* and *Rbpj* dramatically alters gene expression globally (Fig. 5B). Cell annotation based on known marker genes revealed no Müller glia-derived neurons in uninjured and NMDA-treated control samples (Fig. 5D). In contrast, *Nfia/b/x;Rbpj*-deficient MG generated several distinct cell populations, including proliferative Müller glia, neurogenic Müller glia-derived progenitor cells (MGPC), and amacrine and bipolar-like cells, as well as a small number of rod photoreceptors (Fig. 5C). Unexpectedly, we also observed two additional clusters of cells, which selectively expressed *Col25a1* and *Alk*, respectively (Fig. S6A). While the *Col25a1*-positive cluster expressed a subset of photoreceptor/bipolar (*Crx*, *Gnb3, Tacr3*) and RPE (*Rgr*, *Lrat*, *Rrh*)-specific markers, neither of these clearly corresponded to any cell type in wildtype retina. These cells were barely detected in the NMDA-treated retina where substantially higher levels of neurogenesis were observed, implying that they may represent transitional rather than stable cell states (Fig. 5C,D).

**Figure 5:**
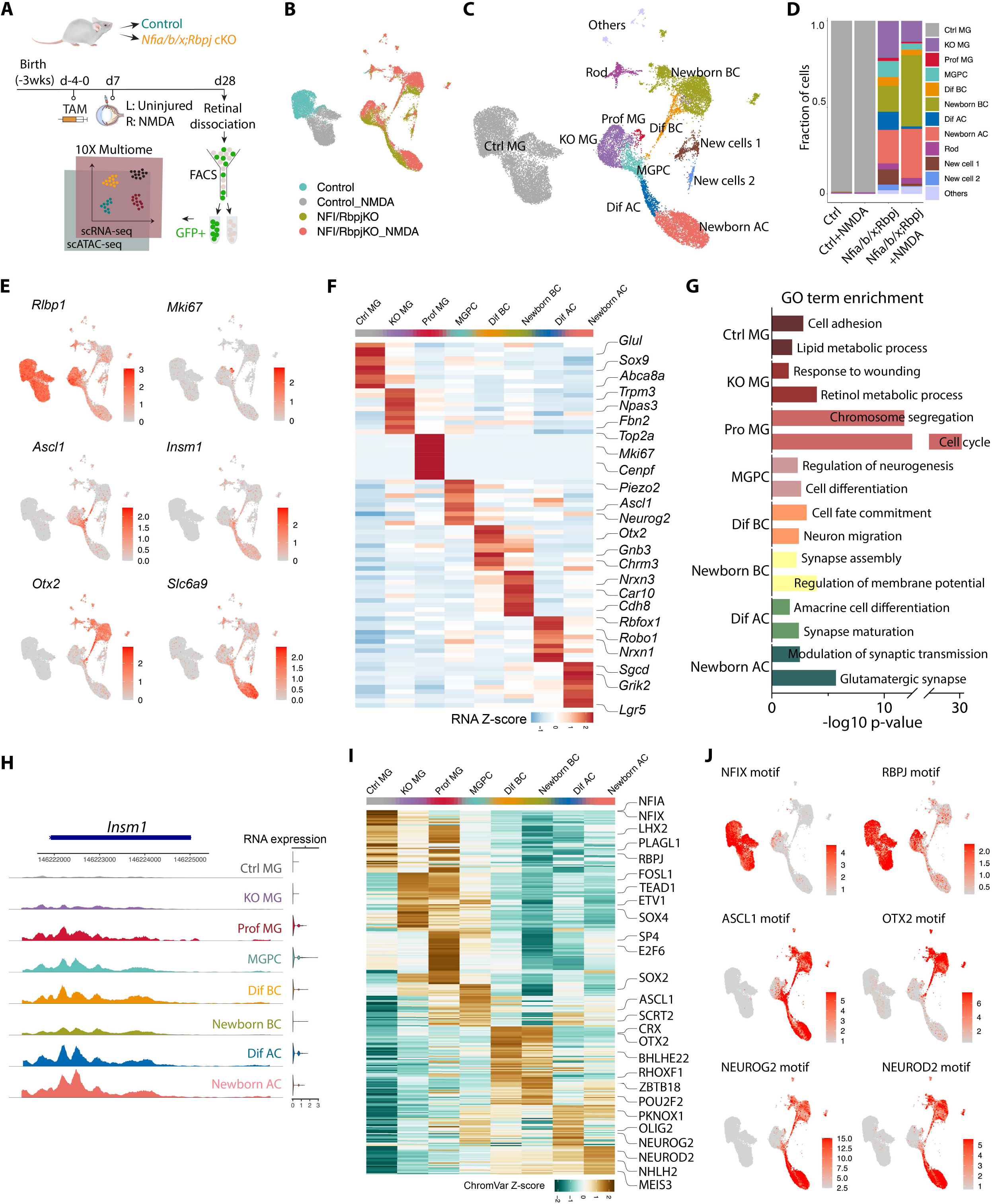
Integrated snRNA/scATAC-Seq analysis of control *and Nfia/b/x*;*Rbpj*-deficient adult Müller glia and their progeny. **(A)** Schematic of the multiomic snRNA/ATAC-seq experimental pipeline. **(B)** UMAP plot of Multiome (scRNA/ATAC-seq) datasets showing the clustering of control and *Nfia/b/x;Rbpj*- deficient MG from uninjured and NMDA-treated retinas. **(C)** UMAP plot showing the identity of cell clusters determined by marker gene expression. **(D)** Stacked barplots represent the proportion of cells in each cluster across different sample groups. **(E)** Feature plots highlighting the cluster of Müller glia (*Rlbp1*), proliferating MG (*Mki67*), neurogenic MGPC (*Ascl1*, *Insm1*), bipolar cells (*Otx2*), and amacrine cells (*Slc6a9*). **(F)** Heatmap showing the expression of top 10 differentially expressed genes (DEGs) for each cell cluster. (**G**) GO enrichment for each cell cluster. The x axis indicates the −log10(p value) of the GO term. **(H)** Representative ATAC peaks and RNA expression level of *Insm1* across different cell clusters. (**I**) Heatmap showing activity of top 50 differential motifs across different cell clusters. (**J**) Feature plots showing activity of indicated motifs. MG, Müller glia; MGPC, Müller glia-derived progenitors; prof MG, proliferating Müller glia, dif BC, differentiating bipolar cells; dif AC, differentiating amacrine cells.

Our multiomic analysis showed that a substantial number of neurons are derived from Müller glia lacking *Nfia/b/x* and *Rbpj* (Fig. 5D). NMDA-induced injury further enhanced the level of Müller glia-derived neurogenesis, which is consistent with our immunohistochemistry data (Fig. 4D, 5D). However, a small subset of Müller glia-derived cells expressed the proliferation marker, *Mki67*, which was undetected in our IHC analysis (Fig. 5E,F). MGPCs selectively expressed neurogenic TFs such as *Ascl1* and *Insm1* (Fig. 5E,G). This MGPC cluster forms clear differentiation trajectories connecting *Rlbp1*-expressing Müller glia to *Otx2/Cabp5*-positive bipolar and *Slc6a9/Elavl3*-positive amacrine cells (Fig. 5E,F, S6A,B). GO enrichment analysis revealed that genes in the proliferating Müller glia cluster were enriched for cell cycle regulation; genes in the MGPC cluster for cell differentiation; and genes in the newborn bipolar and amacrine for membrane potential and synapse regulation (Fig. 5G). While no clear differentiation trajectory connecting neurogenic Müller glia and rod photoreceptors, a subsets of cells in the bipolar differentiation trajectory expressed TFs that promote rod specification (*Prdm1*) or differentiation (*Crx*) ^43–45^, or genes that are known to be selectively expressed in photoreceptor precursors (*Wnt5b*/*Ankrd33b*) ^46^ (Fig. S6B, Table ST3). The origin of the *Col25a1* and *Alk*-positive cell clusters, however, remains less clear.

Many other known cell type-specific genes were selectively upregulated in these cells as they differentiated, including *Car10*/*Lhx4*/*Trpm1* in bipolar cells and *Rbfox3*/*Gad2*/*Neurod2* in amacrine cells (Fig. 5F, Table ST3). These transcriptional changes were reflected in changes in the chromatin accessibility observed in the ATAC-seq analysis. As in *Rbpj*-deficient Müller glia, chromatin accessibility at genes specifically expressed in resting Müller glial genes were decreased, while accessibility for neurogenic TFs such as *Insm1* and *Neurod2* and the rod photoreceptor-specific factor *Nrl* were increased, and maintained in differentiating neurons (Fig. 5H, S6C,D). Analysis of open chromatin regions revealed significant enrichment for motifs for neurogenic TFs such as *Ascl1, Neurog2 and Neurod2,* and reduction in accessibility associated with TF motifs for Müller glia genes – including *Nfia, Nfix, Rbpj*, and *Lhx2* – in the *Nfia/b/x;Rbpj*-deficient Müller glia (Fig. 5I, J). In summary, since no detectable levels of neurogenesis were observed in uninjured *Nfia/b/x*-deficient retina ^6^, and only 5.8% of *Rbpj*-deficient Müller glia undergo conversion to neurons at this age (Fig. 1), these findings indicate that *Nfia/b/x* and *Rbpj* act synergistically to repress neurogenic competence in Müller glia.

### AAV-mediated overexpression of dominant-active Yap5SA promotes proliferation in *Nfia/b/x* and *Rbpj*-deficient retina

While loss of function of *Rbpj* induces Müller glia-derived neurogenesis, we detected no proliferation in *Rbpj*-deficient Müller glia. Moreover, we observed that *Rbpj* loss of function activates expression of cell cycle inhibitors such as *Cdkn1b/c* and *Btg2* (Fig. 3). Likewise, only a limited level of Müller glia proliferation was induced by *Nfia/b/x-* or combined *Nfia/b/x;Rbpj* deletion (Fig. 5). This process of direct transdifferentiation to neurons depletes endogenous Müller glia and severely limits any potential clinical application of Notch pathway inhibition. Recent studies have shown that Hippo signaling represses Müller glia proliferation ^47,48^, and that transgenic mice overexpressing the dominant-active Yap5SA mutant induce Müller glial proliferation^49^. To determine whether overexpression of Yap5SA could induce Müller glia to proliferate, we intravitreally injected adult wildtype control, *Rbpj* and *Nfia/b/x;Rbpj* conditional knockout mice with a Cre-inducible FLEX AAV construct that either stably overexpressed Yap5SA-P2A-mCherry or mCherry alone (Fig. 6A). We then induced Cre activation by 5 daily doses of tamoxifen i.p. injection followed by retinal injury with NMDA. To label proliferating cells, mice were administered EdU via drinking water and daily i.p. injections for 7 days. Retinas were then harvested 3 weeks after NMDA injection for immunostaining to detect Müller glia-derived proliferation and neurogenesis (Fig. 6B).

**Figure 6:**
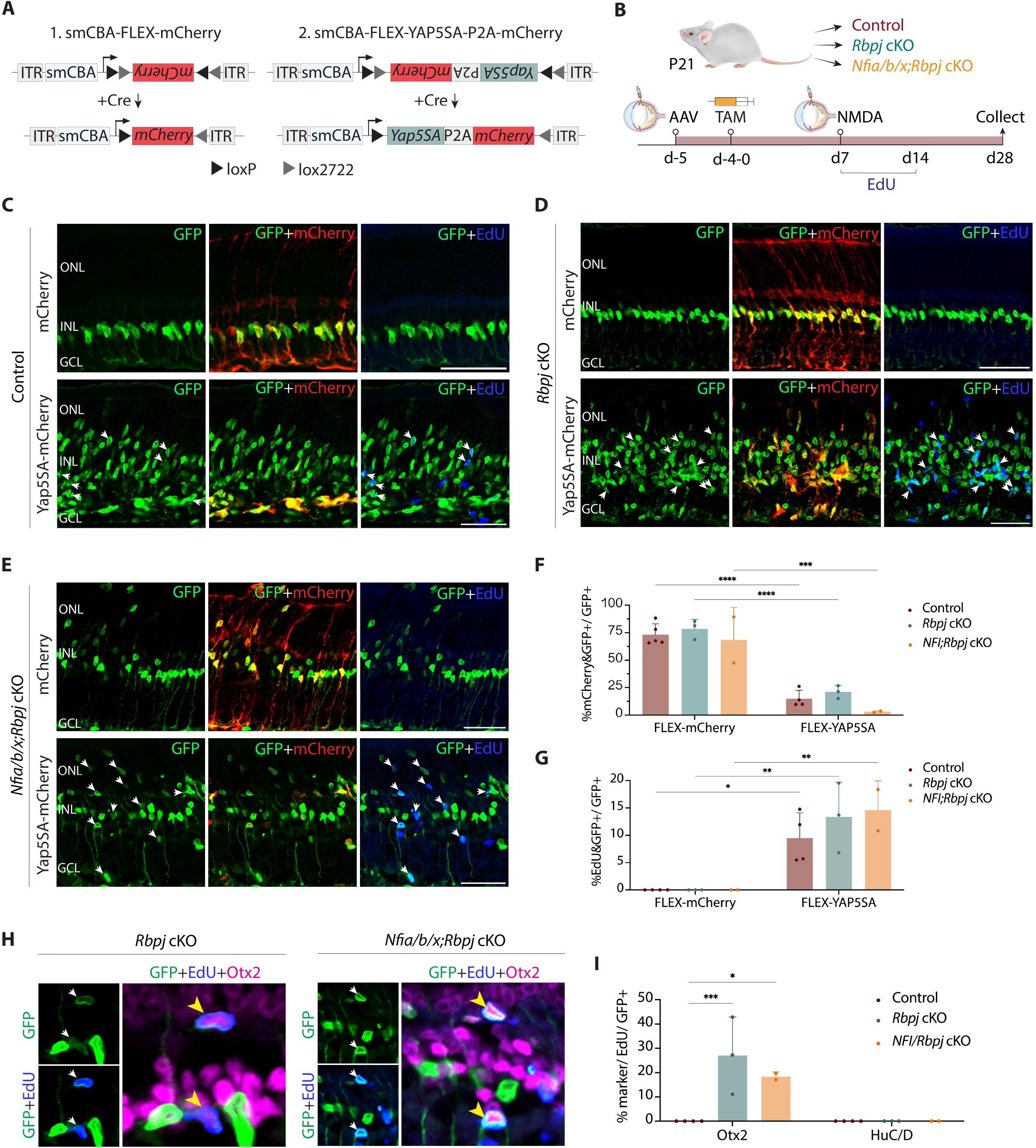
Overexpression of dominant active *YapS5A* stimulates proliferation and enhanced neurogenesis in both *Rbpj* and *Nfia/b/x*+*Rbpj*-deficient adult Müller glia. (A) A schematic of Cre-inducible control and Yap5SA FLEX constructs used in the study. **(B)** Schematic of experimental pipeline. **(C-E)** Representative immunostaining of GFP, mCherry, and EdU on retinal sections from control, *Rbpj* cKO and *Nfia/b/x;Rbpj* cKO with mCherry control and Yap5SA AAVs. White arrowheads indicate co-labeled GFP-positive & EdU-positive cells. **(F)** Quantification of AAV transduction efficiency in MG. **(G)** Quantification of MG co-labeled with EdU. **(H)** Representative immunostaining of EdU-positive MG-derived neurons expressing bipolar cell marker Otx2. Yellow arrowheads indicate EdU/GFP/Otx2 triple-positive cells. **(I)** Quantification of mean percentage ±SD of EdU-positive Müller glia-derived neurons expressing neuronal markers: Otx2 and HuC/D. Significance was determined via two-way ANOVA with Tukey’s test: ****p < 0.0001. Each data point was calculated from an individual retina. TAM, tamoxifen; ONL, outer nuclear layer; INL, inner nuclear layer; GCL, ganglion cell layer. Scale bar = 50μm.

We observed selective Müller glia-specific expression of mCherry in both constructs, as evidenced by colocalization of Sun1-GFP and mCherry (Fig. 6C-E). Little to no EdU/GFP-positive cells were detected in retinas injected with AAV-flex-mCherry, although induction of mCherry is highly efficient (Fig. 6C-F).

Whereas in retinas injected with AAV-flex-Yap5SA-P2A- mCherry, we observed an extensive fraction of GFP/EdU-positive cells across all three sample groups (Fig. 6C-E, G). 9.5% of control, 13.4% of *Rbpj*-, and 14.6% of *Nfia/b/x;Rbpj*-deficient Müller glia colabeled with EdU. We also observed that a significant fraction of EdU/GFP+ cells exhibited faint or no mCherry fluorescent signal. This suggests that YAP5SA overexpression induces Müller glia to undergo multiple cell divisions, subsequently leading to the dilution of AAV mCherry reporter (Fig. 6C-E). In *Nfia/b/x;Rbpj* cKO retinas, which had significant Müller glia depletion (Fig. 4C), we detected an increase in the number of Sox9/GFP+ cells with YAP5SA overexpression (Fig. S7A).

To determine if proliferating Müller glia cells give rise to new neurons, we performed co-immunostaining with neuronal markers Otx2 and HuC/D. While the relative fraction of Otx2/GFP-positive cells was similar between retinas infected with AAV-flex-Yap5SA-P2A-mCherry and AAV-flex-mCherry, we observed that 27.1% and 18.4% of Müller glia-derived cells expressing Otx2 were co-labeled with EdU in *Rbpj-* and *Nfia/b/x;Rbpj*-deficient retinas, respectively. This indicates that they were derived from proliferating Müller glia (Fig. 6H,I). No neurons derived from Müller glia were observed in the control retinas. Interestingly, we did not detect any HuC/D/GFP-positive cells co-labeled with EdU in all three genotypes (Fig. 6I). This observation suggests that proliferating Müller glia induced by YAP5SA overexpression is, for reasons that remain unclear, prevented from differentiating into amacrine cells. Overall, our data revealed that AAV-mediated overexpression of YAP5SA could be used to effectively replenish Müller glia that were depleted due to high levels of non-proliferative transdifferentiation into neurons.

## Discussion

In this study, we identified *Rbpj* and Notch signaling as critical negative regulators of neurogenic competence in mature mammalian Müller glia. Using genetic lineage analysis, immunohistochemistry, and single-cell RNA- and ATAC-Seq, we unambiguously demonstrate that both *Rbpj* and *Notch1/2*-deficient Müller glia cells generate retinal neurons in adult mouse retinas. We showed that loss of function of *Rbpj* induces direct transdifferentiation of Müller glia into bipolar and amacrine cells in the absence of injury, and retinal injury further enhances this effect. We also found that loss of function of *Notch1/2* phenocopies *Rbpj* loss of function. CUT&Tag analysis revealed that Rbpj regulates expression of both Notch pathway components and mature Müller glia-specific genes in Müller glia. Furthermore, we demonstrated that combined deletion of *Rbpj* and *Nfia/b/x* factors resulted in robust levels of Müller glia- derived neurogenesis, indicating that *Nfia/b/x* and *Rbpj* act in parallel pathways to independently inhibit neurogenesis in mature Müller glia. We conducted a comprehensive multiomic analysis to characterize neurogenic glia and newborn neurons. Finally, we showed that AAV-mediated expression of constitutively active Yap strongly induces proliferation and neuron generation in both *Rbpj*-deficient and *Nfia/b/x*;*Rbpj*- deficient Müller glia. These findings further demonstrate that neurogenic competence is actively repressed in mammalian Müller glia, and that disrupting these negative regulators may be of potential therapeutic benefit in developing regenerative therapies for treating retinal dystrophies.

Our findings align with previous studies in zebrafish indicating that Notch signaling maintains Müller glia quiescence and is transiently downregulated following injury ^6,17,19,20^. However, while chemical inhibition of Notch signaling is sufficient to induce Müller glia reprogramming in zebrafish ^14,16^, it is not effective in mice ^16^. Although DAPT-mediated Notch inhibition enhanced neurogenesis in *Nfia/b/x*-deficient Müller glia, this effect was modest compared to the robust effect observed in the *Nfia/b/x;Rbpj* conditional mutant. This discrepancy may be attributed to difficulties in efficiently inhibiting Notch signaling using small molecules, and the fact that *Nfia/b/x* and possibly other factors act in redundant parallel manner to repress neurogenic competence in mammalian Müller glia. Notably, loss of function of both *Rbpj* and *Notch1/2* induces robust levels of Müller glia-derived neurogenesis in the absence of acute injury, in sharp contrast to both *Nfia/b/x* loss of function and *Ascl1* overexpression, where injury is essential for inducing neurogenesis ^5,6^. Only simultaneous overexpression of both *Ascl1* and *Atoh1* has been reported to induce injury-independent Müller glia-derived neurogenesis ^7^. Injury-induced transition to a reactive state is essential for inducing neurogenic competence in zebrafish by downregulating genes that maintain Müller glial quiescence ^6,50^, and these findings imply that inhibition of Notch signaling is an essential component of this process.

During retinal development, a dynamic level of Notch signaling regulates the balance of retinal progenitor maintenance and differentiation of retinal neurons and Müller glia ^51–53, 54^, with Notch pathway inhibition resulting in precocious differentiation of retinal progenitors. Here, we found that inhibition of the Notch pathway led to a direct conversion of Müller glia into retinal interneurons in the absence of detectable proliferation, which contrasts with the limited induction of proliferation seen in *Nfia/b/x*-deficient Müller glia ^6^. This can be reversed by viral misexpression of dominant-active Yap5SA, which induces robust glial proliferation in all genotypes examined, and rescues the loss of Müller glia seen in *Nfia/b/x;Rbpj* conditional mutants. Previous studies that have used constitutive transgenic models to overexpress Yap5SA in Müller glia do not observe tumor mutation or defects in retinal lamination ^55^, implying that excess glia may simply undergo apoptosis and that potential safety concerns may be unexpectedly low.

Moreover, any glial proliferation induced by non-replicative AAV vectors will result in progressive dilution with each round of division, further limiting these risks. However, a full evaluation of the long-term consequences of sustained Yap5SA overexpression in Müller glia remains to be evaluated, as do the effects of sustained Yap activity on differentiation of Müller glia-derived neurons.

While both genetic and chemical inhibition of Notch signaling strongly induce photoreceptor generation in the developing retina ^10,56–58^, we do not detect substantial levels of Müller glia-derived photoreceptor generation in adult *Rbpj* cKO or *Notch1/2* cKO mice. However we do observe limited numbers of Nrl and Recoverin-positive photoreceptor-like cells in the ONL of *Rbpj/Nfia/b/x*-deficient retinas. This relative lack of Müller glia-derived photoreceptors is consistent with previous findings from overexpression of *Ascl1* alone or in combination with other TFs, as well as from *Nfia/b/x*-deficient Müller glia ^5–8^. This finding is surprising, given that over 80% of neurons in the mouse retina are rod photoreceptors and comprise the overwhelming majority of late-born cells. It implies that photoreceptor generation is actively repressed in neurogenic Müller glia, with the underlying reasons remaining unclear. It is likely that a combination of additional treatments that can both promote photoreceptor specification and inhibit these negative regulators will be needed for the development of successful regenerative therapies for photoreceptor dystrophies.

Our findings highlight the importance of targeting active negative regulators of neurogenic competence for effective induction of glia-to-neuron reprogramming. Induced pluripotent stem cells clearly demonstrated the power of TF overexpression as an effective tool for cell conversion ^59^, and most studies aimed at inducing glia-to-neuron conversion in retina ^5,7,8^ and brain ^60,61^ have used similar gain of function approaches. While powerful, high and persistent expression of TFs that promote cell type specification can potentially inhibit terminal differentiation of converted neurons, or generate unwanted neuronal subtypes.

This could potentially be avoided by instead blocking endogenous inhibitors of neurogenesis. Identifying effective methods of doing so, however, has proven challenging. Overexpression of miRNAs which block expression of gliogenic factors can induce fibroblast to neuron conversion *in vitro* ^62–64^, and can enhance the efficiency of glia-to-neuron reprogramming in both brain and retina, but require overexpression of other TFs to induce neurogenesis *in vivo* ^65–68^. Claims of efficient glia-to-neuron conversion in brain and retina by knockdown of the glial-enriched RNA binding protein *Ptbp1* have not been replicated ^22,69–75^, and instead likely reflect technical artifacts resulting from use of AAV-based minipromoter constructs ^76,77^.

Nevertheless, our utilization of rigorous cell lineage analysis and multiome analysis to show that combined loss of function of *Rbpj* and *Nfia/b/x* leads to conversion of over 70% of retinal Müller glia into neurons demonstrates the promise of targeting negative regulatory factors for effective in vivo glia-to-neuron conversion. Since these genes are also expressed in many different glial cell cell types in the brain, this strategy may hold broader potential for developing cell-based therapies for treating neurodegenerative disorders.

## Limitations of the study

Several important questions remain unanswered in this study. First, it is unclear the exact extent to which neurons derived from *Rbpj*- or *Nfia/b/x;Rbpj-*deficient Müller glia resemble endogenous retinal neurons, and whether they are capable of integrating into endogenous retinal circuitry. Second, while their gene expression pattern and morphology closely resemble those of bipolar and amacrine cells, further physiological analysis is essential to fully characterize these neurons. Finally, it remains to be determined whether Müller glia, in which neurogenic competence have been restored by a combination of Notch inhibition and *Nfia/b/x* loss of function, can be guided by additional fate-determining factors to efficiently generate functional photoreceptors or retinal ganglion cells that are lost to blinding diseases.

## Acknowledgements

We thank Dr. Pin Lyu, L. Orzolek, T. Creamer, and members of the Hopkins Single-Cell and Transcriptomic Core for technical assistance. We thank A. Fischer, J. Nathans, and A. Kolodkin, and members of the Blackshaw lab for comments on this manuscript. This work was supported by an award from the National Eye Institute (R01EY031685) to S.B. and a Stein Innovation Award from Research to Prevent Blindness to S.B.

## Data availability

All scRNA-Seq, snRNA-Seq, and scATAC-Seq data described in this study is available as GSE246574.

## Methods

### Mice

Mice were raised and housed in a climate-controlled pathogen-free facility on a 14/10 h light/dark cycle. Mice used in this study were *GlastCreER^T^*^2^*;Sun1-GFP*, which were generated by crossing the *GlastCreER^T^*^2^ and *Sun1-GFP* lines developed by Dr. Jeremy Nathans at Johns Hopkins ^30,78^, and were obtained from his group. *GlastCreER*^T2^;*Rbpj^lox/lox^;Sun1-GFP* mice were generated by crossing *GlastCreER^T^*^2^*;Sun1GFP* with conditional *Rbpj^lox/lox^ mice* (JAX #034200).

*GlastCreER*^T2^;*Notch1^lox/lox^;Notch2^lox/lox^;Sun1-GFP* mice were obtained by crossing *GlastCreER*^T2^;*Notch1^lox/lox^;Sun1-GFP* and *GlastCreER*^T2^;*Notch2^lox/lox^;Sun1-GFP* mice. *GlastCreER*^T2^;*Nfia/b/x^lox/lox^*;*Rbpj^lox/lox^;Sun1-GFP* quadruple cKO mice were generated by crossing *GlastCreER*^T2^;*Nfia/b/x^lox/lox^*;*Sun1-GFP* and *GlastCreER*^T2^;*Rbpj^lox/lox^;Sun1-GFP* mice. Maintenance and experimental procedures performed on mice were in accordance with the protocol approved by the Institutional Animal Care and USe Committee (IACUC) at the Johns Hopkins School of Medicine.

### Intraperitoneal tamoxifen (TAM) Injection

To induce *Cre* recombination, animals at ∼3 weeks of age were intraperitoneally injected with tamoxifen (Sigma-Aldrich, #H6278-50mg) in corn oil (Sigma-Aldrich, #C8267-500ML) at 1.5mg/dose for five consecutive days.

### NMDA treatment

Adult mice were anesthetized with isoflurane inhalation. Two microliters of 100 mM NMDA in PBS were intravitreally injected using a microsyringe with a 33G blunt-ended needle.

### DAPT treatment

Adult *Glast^CreERT^*^2^*;Nfia/b/x^lox/lox^; Sun1-GFP* mice were anesthetized with isoflurane inhalation and intravitreally injected with 200 μM (2 μL/injection) of DAPT (MedChemExpress, #HY-13027) in DMSO using a microsyringe with a 33G blunt-ended needle.

### Cloning and Production of Adeno-Associated Virus

The Addgene #50462 construct containing an EF1a promoter was replaced with a smCBA promoter. The P2A ribosomal self-cleaving peptide is used to simultaneously express Yap5SA 5’ to the mCherry reporter as a single polypeptide, which is then cleaved to generate the transcription factor and mCherry. The coding sequence of Yap5SA was synthesized by GeneWiz. AAV constructs were packaged into AAV2.7m8 by Boston Children’s Hospital Viral Core.

### FLEX-AAV Delivery

∼3 weeks old old *GlastCreER^T^*^2^*;Sun1-GFP* and *GlastCreER*^T2^;*Rbpj^lox/lox^;Sun1-GFP* mice were anesthetized with isoflurane inhalation and intravitreally injected with AAV-FLEX constructs using a microsyringe with a 33G blunt-ended needle. One day following AAV transduction, 5 daily doses of TAM (1.5mg/dose in corn oil) were administered to activate Cre recombinase. Titre and injection volume for each construct are listed below:

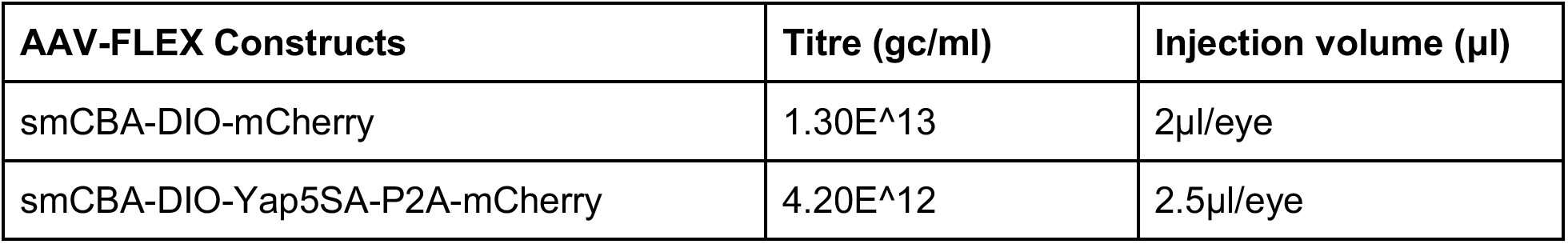

### Immunohistochemistry and imaging

Collection and immunohistochemical analysis of retinas were performed as described previously (Hoang et al., 2020). Briefly, mouse eye globes were fixed in 4% paraformaldehyde (ElectronMicroscopySciences, #15710) for 4 h at 4°C. Retinas were dissected in 1x PBS and incubated in 30% sucrose overnight at 4°C.

Retinas were then embedded in OCT (VWR, #95057-838), cryosectioned at 16 μm thickness, and stored at −20°C. Sections were dried for 30 min in a 37°C incubator and washed 3 × 5 min with 0.1% TritonX-100 in PBS (PBST). EdU labeling was performed by using Click-iT EdU kit (ThermoFisher, #C10340, #C10636) following the manufacturer’s instructions. Sections were then incubated in 10% Horse Serum (ThermoFisher, #26050070), 0.4% TritonX-100 in 1x PBS (blocking buffer) for 2 h at room temperature (RT) and then incubated with primary antibodies in the blocking buffer overnight at 4°C.

Sections were washed 4 x 5 min with PBST to remove excess primary antibodies and were incubated in secondary antibodies in blocking buffer for 2 h at RT. Sections were then counterstained with DAPI in PBST, washed 4 x 5 min in PBST and mounted with ProLong Gold Antifade Mountant (Invitrogen, #P36935) under coverslips (VWR, #,48404-453), air-dried, and stored at 4°C. Fluorescent images were captured using a Zeiss LSM 700 confocal microscope.

### Cell quantification and statistical analysis

Otx2/GFP-positive and HuC/D/GFP-positive cells were counted and divided by the total number of GFP- positive cells from a single random whole section per retina. For cell proliferation quantification, EdU/GFP- positive cells were counted and divided by the total number of GFP-positive cells from a single random whole section per retina. EdU-positive MG-derived neurons were quantified by the number of EdU/GFP/Otx2-positive cells divided by the total GFP/Otx2-positive cells. AAV infection efficiency were calculated by the number of mCherry/GFP-positive cells divided by the total number of GFP-positive cells. Each data point in the bar graphs was calculated from an individual retina. All cell quantification data were graphed and analyzed using GraphPad Prism 10. Two-way ANOVA were used for analysis between 3 or more samples of multiple groups. All results are presented as mean ± SD.

### Retinal cell dissociation

Retinas were dissected in fresh ice-cold PBS and retinal cells were dissociated using an optimized protocol as previously described ^79^. Each sample contains a minimum of 4 retinas from 4 animals of both sex.

Dissociated cells were resuspended in ice-cold HBAG Buffer containing Hibernate A (BrainBits, #HALF500), B-27 supplement (ThermoFisher, #17504044), and Glutamax (ThermoFisher, #35050061).

### CUT&Tag library preparation

CUT&Tag was performed using the CUTANA CUT&Tag kit following the manufacturer’s instructions. Adult *GlastCreER^T^*^2^*;Sun1-GFP* mice were induced with tamoxifen injection 1 week prior to retinal dissociation for CUT&Tag experiment. Nuclei were isolated from ∼100,000 FACS-isolated GFP-positive cells in NE Buffer for 10 min on ice. Primary antibodies used were rabbit anti-IgG (negative control), rabbit anti-H3K27me3 (positive control), and rabbit anti-Rbpj. Libraries were sequenced on NextSeq Mid150 with 130 million reads per library.

### CUT&Tag data analysis

To analyze the CUT&Tag data, Fastq files were trimmed for adapters by Cutadapt and mapped to the mm10 genome by Bowtie2 ^80^ using the parameters: --local --very-sensitive -no-mixed --no-discordant -I 10 - X 700. Bam files were generated and multimapping reads were removed by SAMtools ^81^, and duplicate reads were removed by Picard. Pileups were generated as bigwig files using deepTools ^82^ function ‘bamCoverage’ with counts per million (CPM) normalization and visualized in IGV. Replicates from each biological group i.e., IgG, H3K4me3, Rpbj, were merged prior to peak calling using SAMtools. MACS2^83^ was used to call peaks following a standard procedure. The heatmaps showing peak distribution for each merged dataset of IgG, H3K4me3, and Rbpj were generated using the deepTools function ‘plotHeatmap’. Motif-enrichment analysis was performed using MEME function ‘Enriched motif’ ^84^; only the top 5000 peaks ranked by the p-adjusted values on the merged data for each group were used for this analysis.

### Single-cell RNA-sequencing (scRNA-seq) library preparation

scRNA-seq were prepared on dissociated retinal cells using the 10X Genomics Chromium Single Cell 3’ Reagents Kit v3.1 (10X Genomics, Pleasanton, CA). Libraries were constructed following the manufacturer’s instructions and were sequenced using Illumina NextSeq. Sequencing data were processed through the Cell Ranger 7.0.1 pipeline (10X Genomics) using default parameters.

### Single-cell ATAC-sequencing (scATAC-seq) library preparation

scATAC-seq were prepared on FACS-isolated cells using the 10X Genomics Chromium NextGEM SingleCell ATAC Reagent Kits v1.1 (10X Genomics, Pleasanton, CA). Briefly, cells were spun down at 500xg for 5 min and resuspended in 100 μl of ice-cold 0.1x Lysis Buffer and lysed by pipette-mixing 4 times and incubated on ice for 4 min total. Cells were washed with 0.5 ml of ice-cold Wash Buffer and spun down at 500xg for 5 min at 4°C. Nuclei pellets were resuspended in 10-15 μl Nuclei Buffer and counted using Trypan blue. Resuspended cell nuclei (10-15k) were used for transposition and loaded into the 10x Genomics Chromium Single Cell system. ATAC libraries were amplified with 10 PCR cycles and were sequenced on Illumina NovaSeq with ∼200 million reads per library. Sequencing data were processed through the Cell Ranger ATAC 1.1.0 pipeline (10X Genomics) using default parameters.

### Single-cell Multiome ATAC + GEX sequencing library preparation

scATAC-seq and scRNA-seq were prepared on FACS-isolated GFP-positive cells using the 10X Genomic Chromium Next GEM Single Cell Multiome ATAC + Gene Expression kit following the manufacturer’s instructions. Briefly, cells were spun down at 500xg for 5 min and resuspended in 100 μl of ice-cold 0.1x Lysis Buffer and lysed by pipette-mixing 4 times and incucated on ice for 4 min total. Cells were washed with 0.5 ml of ice-cold Wash Buffer and spun down at 500xg for 5 min at 4°C. Nuclei pellets were resuspended in 10-15 μl Nuclei Buffer and counted using Trypan blue. Resuspended cell nuclei (10-15k) were used for transposition and loaded into the 10x Genomics Chromium Single Cell system. ATAC libraries were amplified with 10 PCR cycles and were sequenced on Illumina NovaSeq with ∼200 million reads per library. RNA libraries were amplified from cDNA with 14 PCR cycles and were sequenced on Illumina NovaSeq 6000.

### scRNA-seq data analysis

scRNA-seq of the control and *Rbpj* cKO mice were analyzed using the Seurat package^85^. Briefly, the gene expression data were jointly normalized by function ‘NormalizeData’, the PCA dimensionality reduction, Louvain clustering, and UMAP visualization were performed on the top 30 principal components. Müller glia (MG) cells then were subset from other cell types by the function ‘subset’, and dimensionality reduction and visualization were performed a second time on the top 30 principal components. Expression of key marker genes within each cell cluster were used to confirm the appropriate assignment of cell types.

### scATAC-seq data analysis

scATAC-seq of the control and Rbpj KO mice were analyzed using the Signac package^86^. Briefly, MG were subset from other cell types, dimensionality reduction was performed using functions ‘FindClusters’ and ‘FindNeighbors’. UMAP visualization was performed on the top 30 principal components. Enhanced accessibility of marker gene bodies and enrichment of marker TF motifs within each MG cell were used to confirm the appropriate assignments. The scATAC-seq peak calling was performed using MACS2 and the differential accessibility regions (DARs) were obtained using function ‘FindMarker’ comparing between the control and *Rbpj cKO* groups. The high-confidence peaks were used for chromatin accessibility heatmap and depth-normalized pileups from MACS2 were used for genome-browser visualization in IGV^87^. The scATAC-seq motif enrichment was performed using the function ‘chromVAR’. The footprinting information for sets of motifs was obtained and visualized using the Signac functions ‘Footprint’ and ‘PlotFootprint’, respectively.

### Multiomic data analysis

For the data processing, raw scRNA- and scATAC-seq data were processed with the Cell Ranger software for formatting reads, demultiplexing samples, genomic alignment, and generating the cell-by-gene count matrix. The cell-by-gene count matrix is the final output from the Cell Ranger pipeline and was used for all downstream analysis. Then, Seurat ^85^ and Signac ^86^ packages were used to create Seurat objects for each sample with the cell-by-gene count matrix with the function ‘CreateSeuratObject’. After visual checking the violin plot of the total counts for each cell, cells with nCount_ATAC > 100,000 & nCount_RNA > 30,000 & nCount_ATAC < 1,000 & nCount_RNA < 1,000 & nFeature_RNA < 500 & nucleosome_signal > 1.2 & TSS.enrichment < 2 were filtered out. Additionally, cells with a mitochondrial fraction > 15% were also removed.

For the data analysis, the weighted nearest neighbor (WNN) method was used to jointly analyze a single-cell dataset measuring both DNA accessibility and gene expression in the same cells using Signac and Seurat function ‘FindMultiModalNeighbors’ ^88^. The ‘SCTransform’ and ‘RunTFIDF’ functions were used to normalize the data for the scRNA-seq and scATAC-seq analysis, respectively. The 2D UMAP was generated using the first 30 dimensions for both the scRNA-seq and scATAC-seq. Cluster-based differential gene expression (DEGs) and differential accessibility regions (DARs) between cell types were performed using the Seurat and Signac function ‘FindAllMarkers’. The average gene expression and peak of each cell type were obtained using the Seurat and Signac function ‘AverageExpression’. The transcription factor motif activity per cell was performed using Signac function ‘RunChromVAR’’. The average gene expression, average peak, and average motif score were visualized in the heatmaps using Seurat and Signac function ‘Doheatmap’.

### Gene Ontology (GO) analysis

To obtain the significantly enriched Gene Ontology terms (biological process), the Database for Annotation Visualization and Integrated Discovery (DAVID) ^89^ was applied under the cut-off of p-value < 0.01.

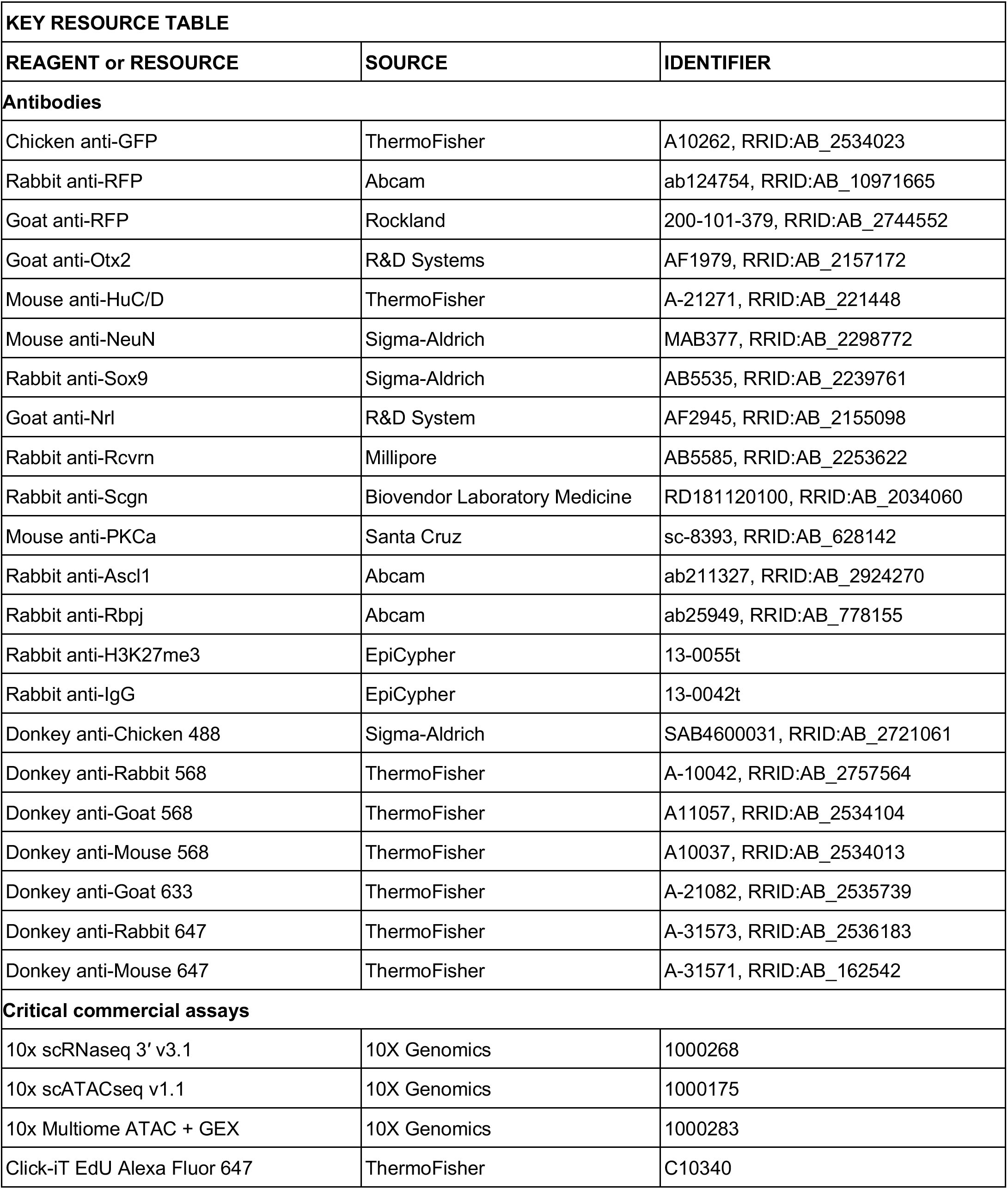
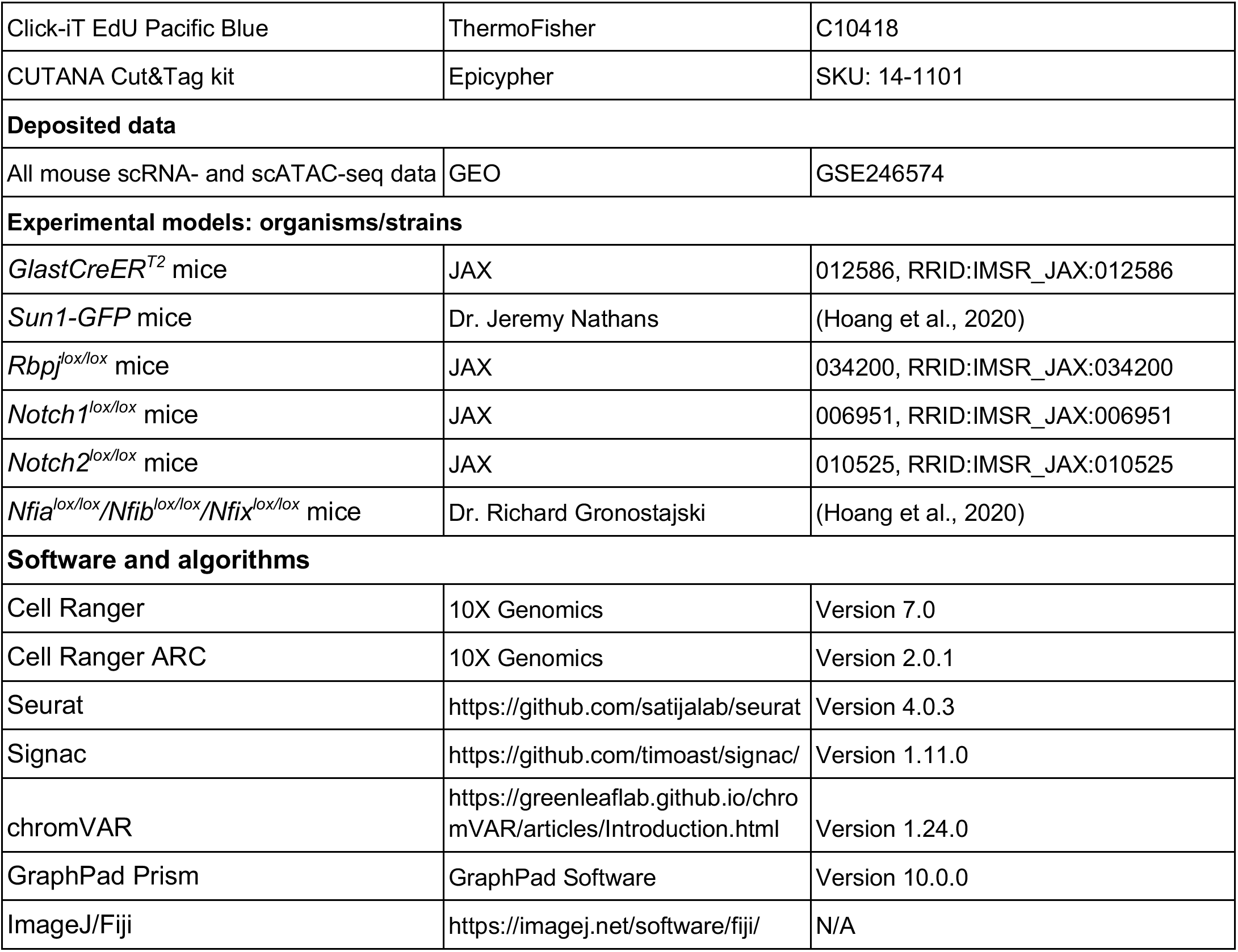

---

**Figure S1:**
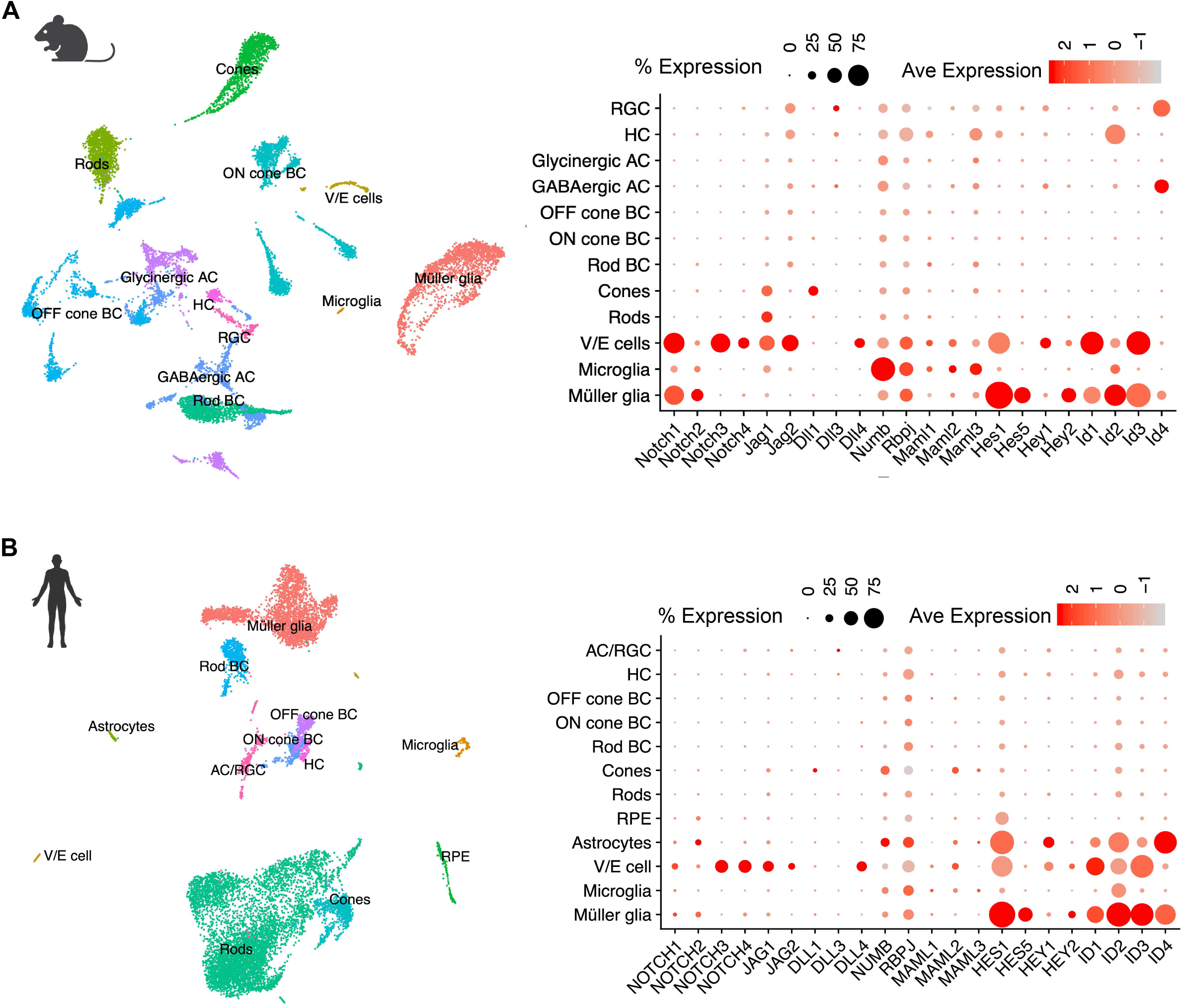
ScRNA expression of Notch signaling components in adult mouse and human retinas. **(A)** UMAP plot of scRNA-Seq data from adult mouse retina from ^22^, and Dot Plot showing cellular expression levels of Notch pathway components in each major cell type. **(B)** UMAP of scRNA-Seq data from adult human retina from ^21^, and Dot Plot showing cellular expression levels of Notch pathway components in each major cell type.

**Figure S2:**
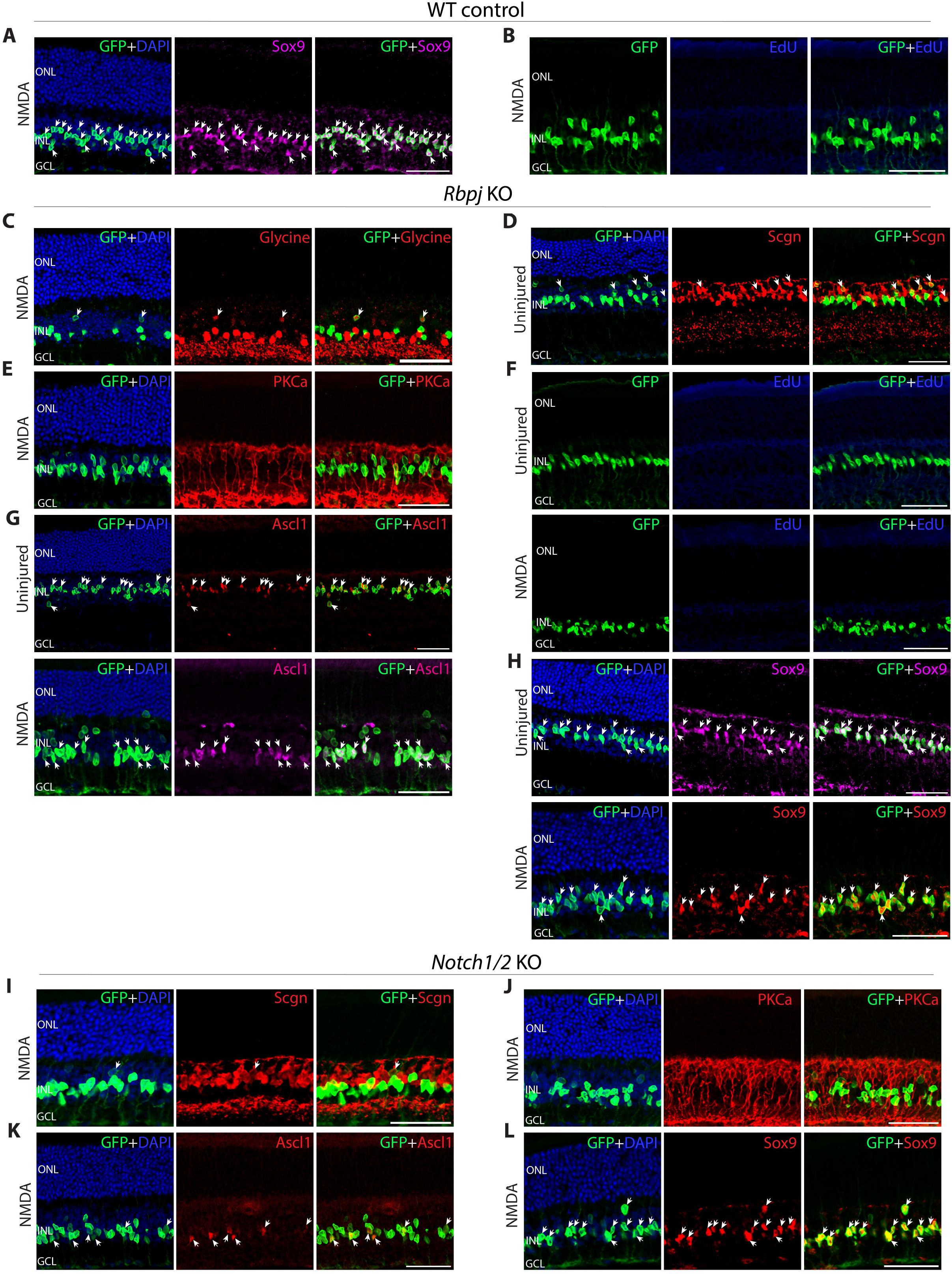
Immunohistochemical analysis of glial and neuronal markers and EdU incorporation following loss of function of *Rbpj* or *Notch1/2* in adult Müller glia. (**A**,**B**) Representative images of control retina immunolabeled for GFP and **(A)** Sox9, **(B)** EdU. (**C**- **H**)Representative images of *Rbpj*-deficient retina immunolabeled for GFP and **(C)** Glycine, **(D)** Scgn, **(E)** PKCa, **(F)** EdU, **(G)** Ascl1, **(H)** Sox9. (**I**-**L**) Representative images of *Notch1/2*-deficient retina immunolabeled for GFP and **(I)** Scgn, **(K)** PKCa, **(K)** Ascl1, **(L)** Sox9. White arrowheads indicate cells colabeled with both GFP and the relevant marker. ONL, outer nuclear layer; INL, inner nuclear layer; GCL, ganglion cell layer. Scale bar = 50μm.

**Figure S3:**
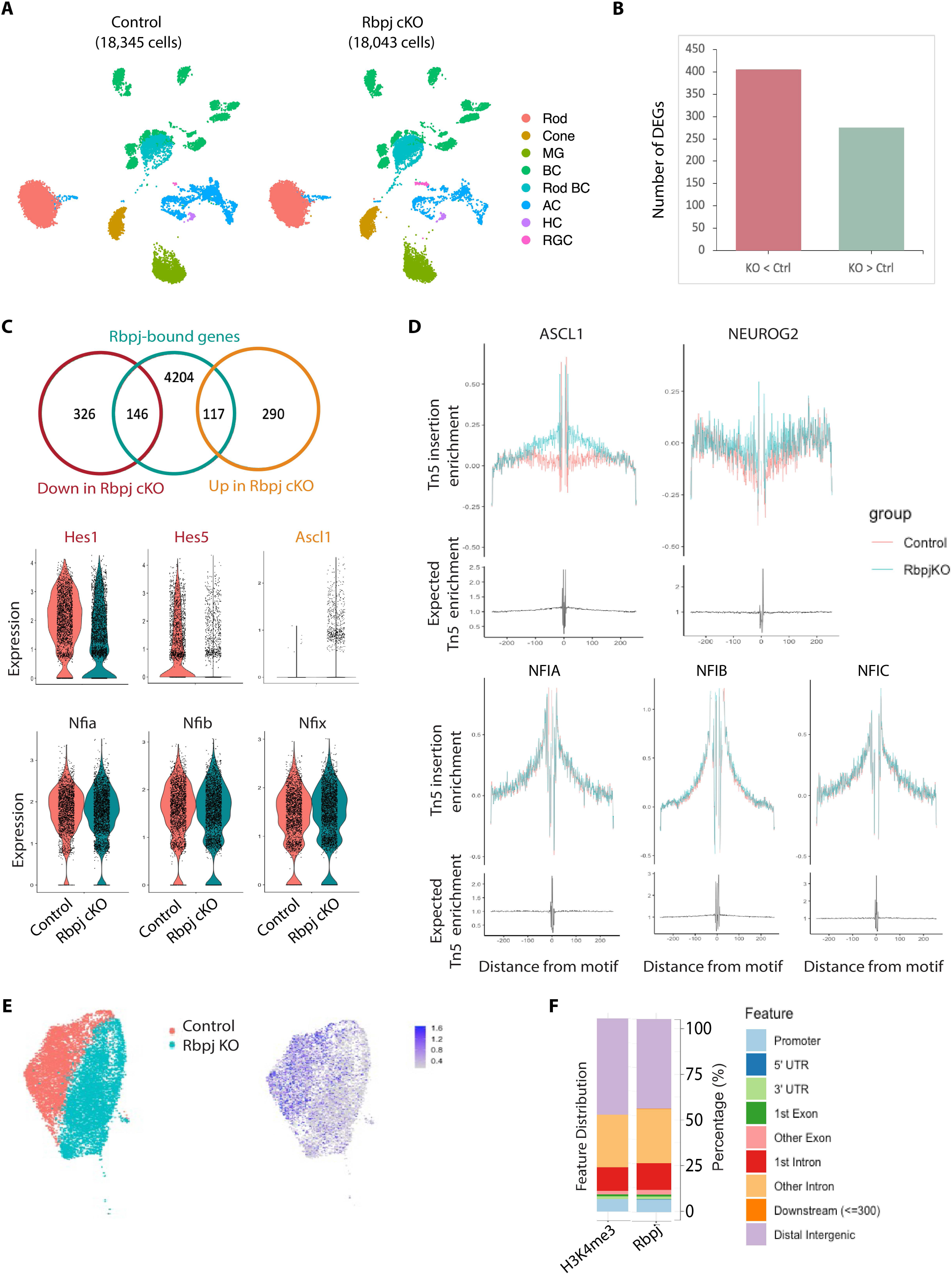
Integrated multiomic analysis to identify Rbpj target genes. **(A)** scRNA-Seq clustering of control and *Rbpj* cKO uninjured whole retinas at 1 week after tamoxifen injection. **(B)** Number of differentially expressed genes between control and *Rbpj*-deficient Müller glia. **(C)** Integrative analysis of CUT&Tag and scRNA-seq data to identify genes regulated by Rbpj. Violin plots showing gene expression levels of *Hes1/5, Ascl1, Nfia/b/x* in control and *Rbpj*-deficient MG. **(D)** TF footprint profiles for *Ascl1*, *Neurog2*, and *Nfia/b/x*. Tn5 insertion tracks are shown below. **(E)** Clustering of scATAC-seq data showing a reduction of Rbpj motif activity in the *Rbpj*-deficient Müller glia, as expected. **(F)** Distribution of Cut&Tag sequencing reads from H3K4me3 and Rbpj samples mapping to genomic features.

**Figure S4:**
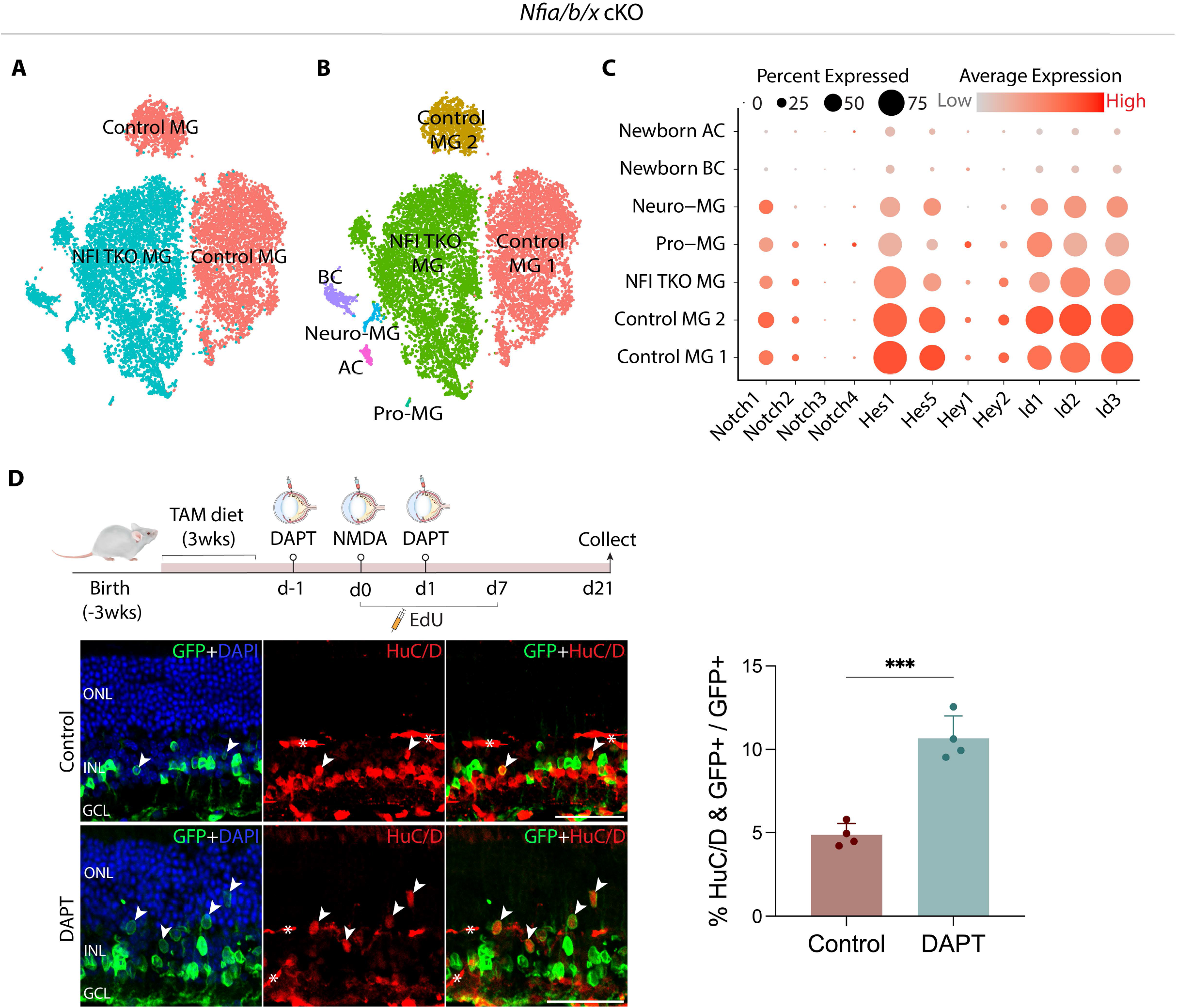
Notch signaling is retained in *Nfia/b/x*-deficient Müller glia, and DAPT-mediated Notch inhibition enhances reprogramming. **(A)** Clustering of control and *Nfia/b/x*-deficient MG **(B)** Cell clusters determined by marker gene expression. **(C)** Dot plot showing gene expression and cell percentages for Notch pathway genes for different cell populations. **(D)** Representative IHC for GFP and HuC/D expression in control and DAPT-treated retinas. White arrowheads indicate colabeled GFP+/marker+ cell. *Asterisks indicate mouse-on-mouse vascular staining. Quantification of mean percentage ±SD HuC/D/GFP+ / GFP+ cells. Significance was determined via unpaired *t* test: ****p < 0.0001. *n=4*. ONL, outer nuclear layer; INL, inner nuclear layer; GCL, ganglion cell layer. Scale bar = 50 μm.

**Figure S5:**
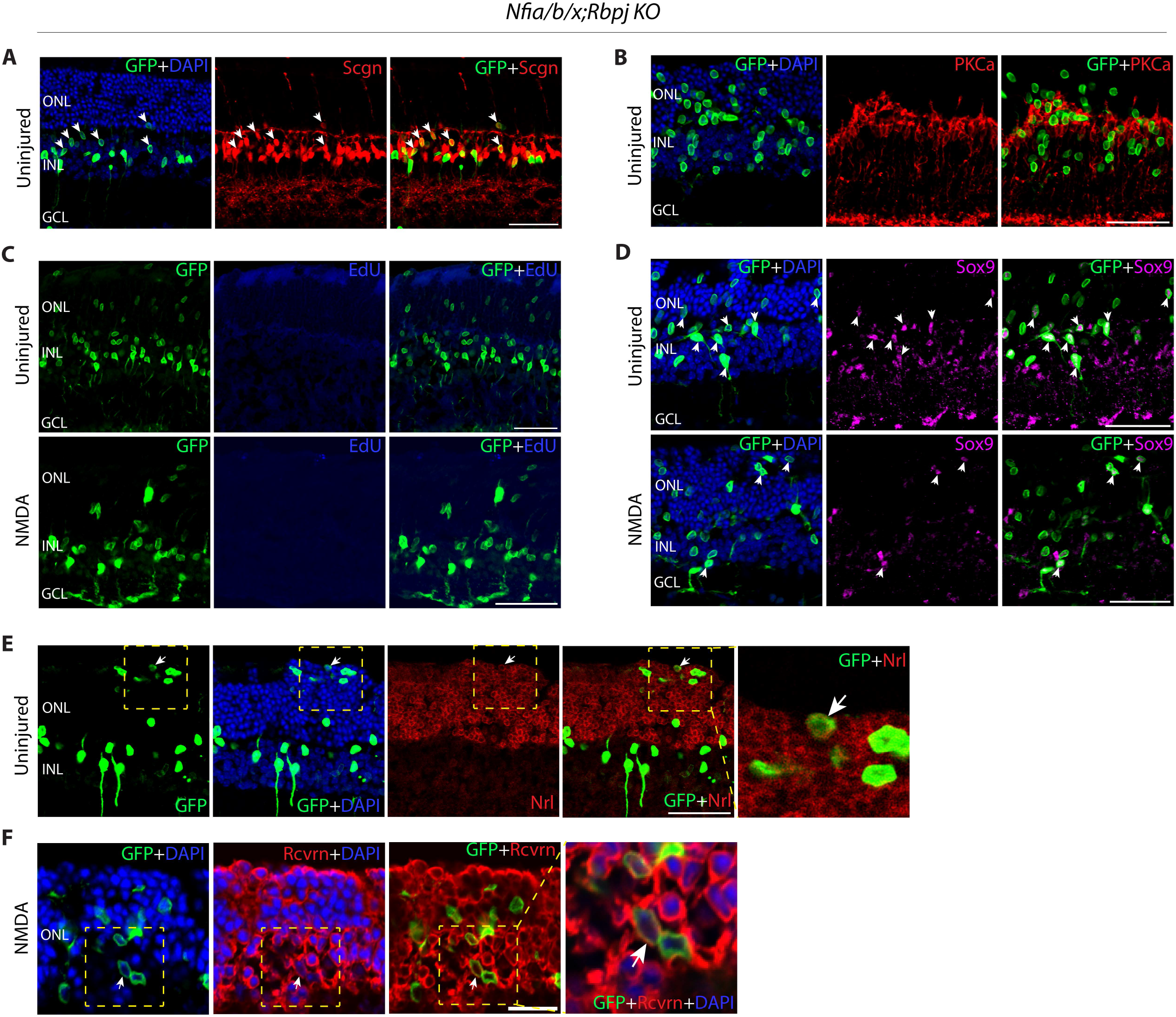
Immunohistochemical analysis of additional markers and EdU incorporation following loss of function of *Nfia/b/x* and *Rbpj* in adult Müller glia. Representative images of retinal sections immunolabeled for GFP and **(A)** Scgn, **(B)** PKCa, **(C)** EdU, **(D)** Sox9, **(E)** Nrl, and **(F)** Recoverin. White arrowheads indicate cells colabeled with both GFP and the relevant marker. ONL, outer nuclear layer; INL, inner nuclear layer; GCL, ganglion cell layer. Scale bar = 50μm (A-E) and 20μm (F).

**Figure S6:**
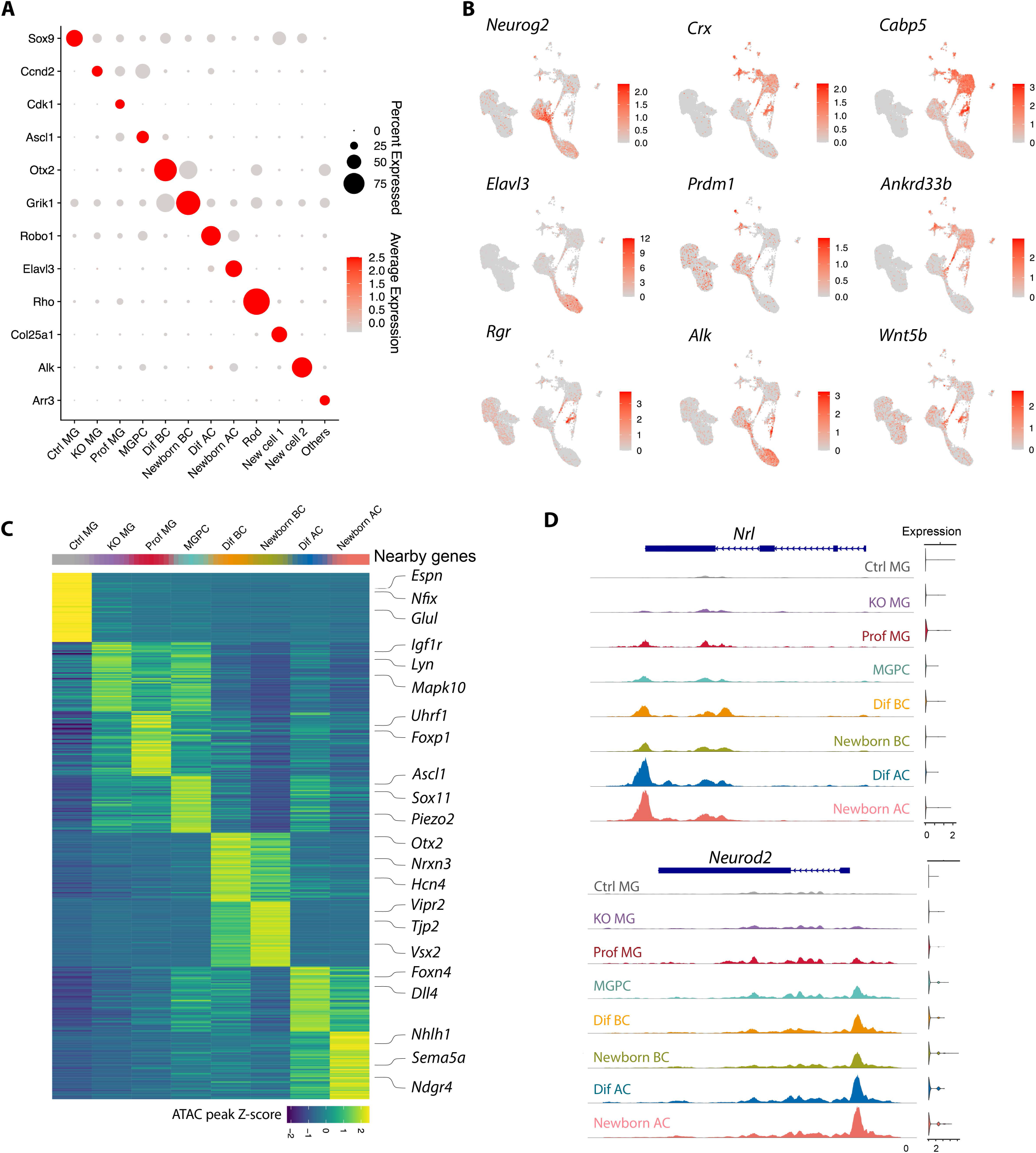
Additional analysis of multiomic data from wildtype control and *Nfia/b/x;Rbpj*-deficient retinas. **(A)** Dot plot showing gene expression and cell percentages for Müller glia, neurogenic and neuronal marker genes for different cell populations. **(B)** Feature plots highlighting the cluster of MGPC (*Neurog2*), bipolar cell (*Cabp5*), amacrine cell (*Elavl3*), photoreceptors and their precursors (*Prdm1, Ankrd33b, Wnt5b, Arr3, Crx, Rgr*). **(C)** Heatmap of top 50 differential ATAC peaks for each cell population and selected nearby genes. **(D)** Representative ATAC peaks and RNA expression levels for *Nrl* and *Neurod2* across different cell clusters.

**Figure S7:**
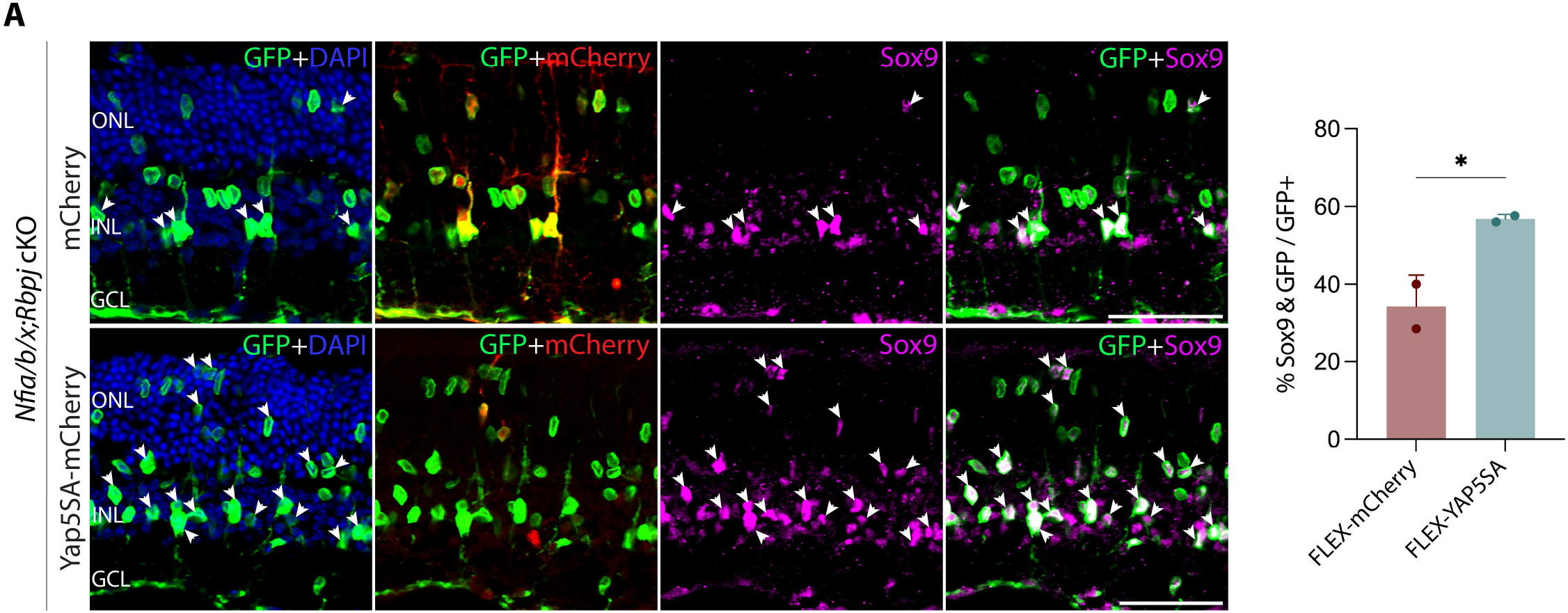
Increased Sox9-positive *Nfia/b/x;Rbpj*-deficient Müller glia following AAV-mediated overexpression of YAP5SA. **(A)** Representative immunostaining for GFP, mCherry and Sox9 in *Nfia/b/x;Rbpj* cKO retinas. White arrowheads indicate co-labeled GFP/Sox9+ cells. Quantification of mean percentage ±SD Sox9/GFP+ cells. Significance was determined via unpaired *t* test: ****p < 0.0001. *n=2*. ONL, outer nuclear layer; INL, inner nuclear layer; GCL, ganglion cell layer. Scale bar = 50 μm.

## Supplemental tables

**Table ST1:** List of genes, chromatin regions, and transcription factor motifs that are differentially regulated in *Rbpj*-deficient Müller glia+MGPCs (scRNA-Seq and scATAC-Seq).

**Table ST2:** List of genes and genomic loci directly targeted by RBPJ in Müller glia.

**Table ST3:** List of genes, chromatin regions, and transcription factor motifs that are differentially regulated in wildtype control, *Nfia/b/x+Rbpj*-deficient Müller glia and their progenies.

---

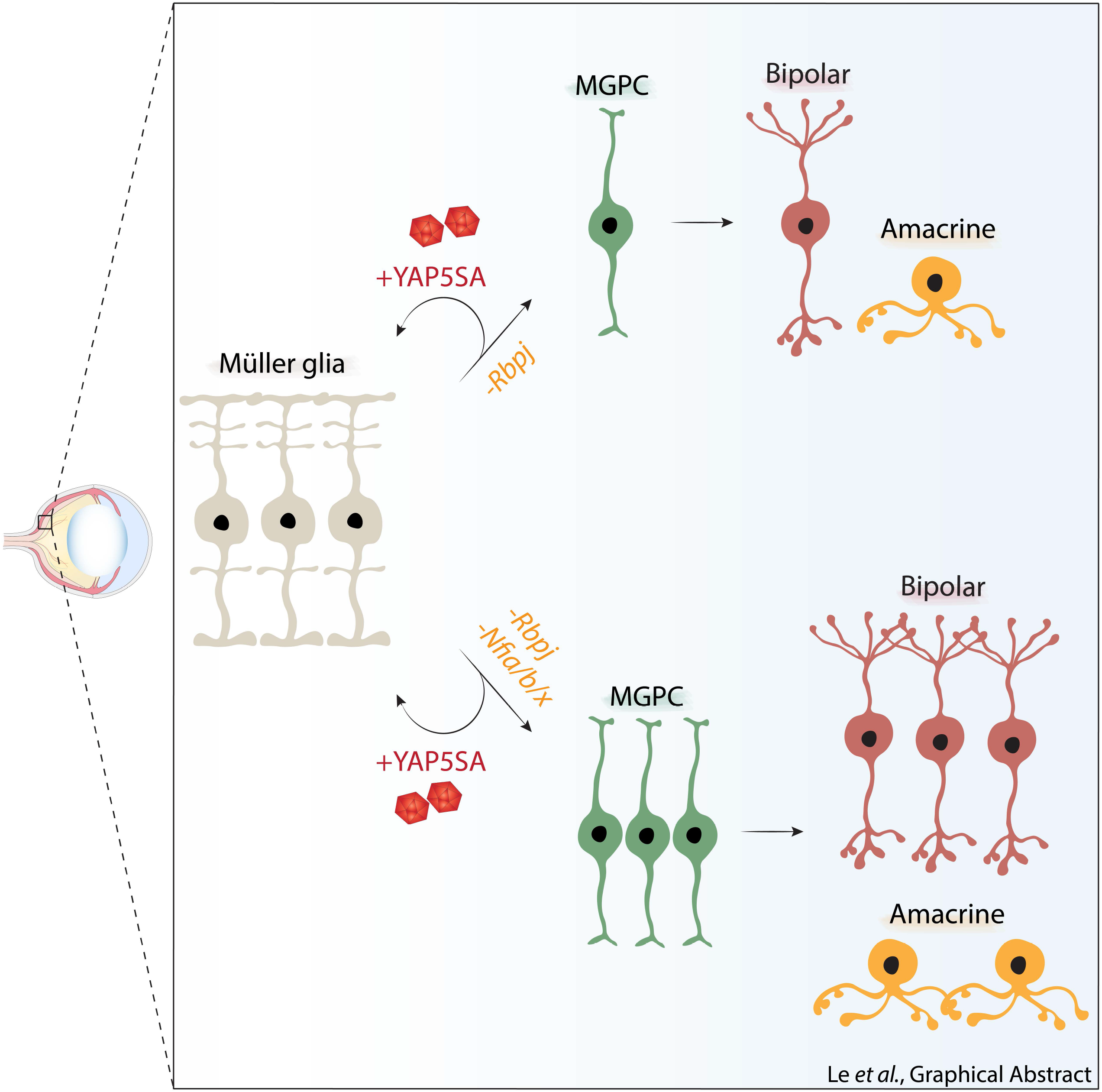

